# Predictive Network Analysis Identifies *JMJD6* and Other Novel Key Drivers in Alzheimer’s Disease

**DOI:** 10.1101/2022.10.19.512949

**Authors:** Julie P. Merchant, Kuixi Zhu, Marc Y.R. Henrion, Syed S.A. Zaidi, Lau Branden, Sara Moein, Melissa L. Alamprese, Richard V. Pearse, David A. Bennett, Nilüfer Ertekin-Taner, Tracy L. Young-Pearse, Rui Chang

**Affiliations:** Ann Romney Center for Neurologic Diseases, Brigham and Women’s Hospital and Harvard Medical School, Boston, MA, USA; The Center for Innovation in Brain Sciences, University of Arizona, Tucson, AZ, USA; Liverpool School of Tropical Medicine, Pembroke Place, Liverpool, Pembroke Place, L3 5QA, UK; Malawi - Liverpool - Wellcome Trust Clinical Research Programme, PO Box 30096, Blantyre, Malawi; Arizona Research Labs, Genetics Core, University of Arizona, Tucson, AZ, USA; Rush Alzheimer’s Disease Center, Rush University Medical Center, Chicago, IL, USA; Department of Neuroscience, Mayo Clinic Florida, Jacksonville, FL, USA; Department of Neurology, Mayo Clinic Florida, Jacksonville, FL, USA; Harvard Stem Cell Institute, Harvard University, Boston, MA, USA; Department of Neurology, University of Arizona, Tucson, AZ, USA; INTelico Therapeutics LLC, Tucson, AZ, USA; PATH Biotech LLC, Tucson, AZ, USA

**Author notes:** Authors equally contributed to the manuscript (Julie P. Merchant, Kuixi Zhu).

## Abstract

Despite decades of genetic studies on late onset Alzheimer’s disease (LOAD), the molecular mechanisms of Alzheimer’s disease (AD) remain unclear. Furthermore, different cell types in the central nervous system (CNS) play distinct roles in the onset and progression of AD pathology. To better comprehend the complex etiology of AD, we used an integrative approach to build robust predictive (causal) network models which were cross-validated over multiple large human multi-omics datasets in AD. We employed a published method to delineate bulk-tissue gene expression into single cell-type gene expression and integrated clinical and pathologic traits of AD, single nucleotide variation, and deconvoluted gene expression for the construction of predictive network models for each cell type in AD. With these predictive causal models, we are able to identify and prioritize robust key drivers of the AD-associated network state. In this study, we focused on neuron-specific network models and prioritized 19 predicted key drivers modulating AD pathology. These targets were validated via shRNA knockdown in human induced pluripotent stem cell (iPSC) derived neurons (iNs), in which 10 out of the 19 neuron-related targets (*JMJD6, NSF, NUDT2, YWHAZ, RBM4, DCAF12, NDRG4, STXBP1, ATP1B1*, and *FIBP*) significantly modulated levels of amyloid-beta and/or phosphorylated tau peptides in the postmitotic iNs. Most notably, knockdown of *JMJD6* significantly altered the neurotoxic ratios of Aβ42 to 40 and p231-tau to total tau, indicating its potential therapeutic relevance to both amyloid and tau pathology in AD. Molecular validation by RNA sequencing (RNAseq) in iNs further confirmed the network structure, showing significant enrichment in differentially expressed genes after knockdown of the validated targets. Interestingly, our network model predicts that these 10 key drivers are upstream regulators of REST and VGF, two recently identified key regulators of AD pathogenesis.

## Introduction

Late-onset Alzheimer’s Disease (LOAD) is the leading cause of dementia, which is characterized by progressive impairments in memory, cognition, and executive functions, along with behavioral and psychiatric symptoms including agitation, aggression, mood disorders, and psychosis[1]. The hallmark features of AD include pathological aggregation of extracellular plaques, composed of amyloid-β (Aβ) peptides, and intracellular neurofibrillary tangles (NFTs), composed of hyperphosphorylated tau (p-tau) protein[2], which lead to neuron death. Genome-wide association studies (GWAS) have implicated over 30 loci associated with AD risk[3–16]. In previous studies, we and others have shown that LOAD is a complex pathological process involving an interactive network of pathways among multiple cell types in the brain (neurons, microglia, astrocytes, etc.) influenced by genetic variation, aging, and environmental factors[17, 18]. Implicated pathways include those involved in mitochondrial metabolism, response to unfolded proteins, immune response, phagocytosis, and synaptic transmission[19–22]. The complexity of these multi-modal (cell type) networks highlights the necessity to study networks of molecular interactions by cell type and to identify cell-type specific pathways and key drivers in AD. In this study, we developed a multi-step pipeline using advanced computational systems biology approaches to construct robust data-driven neuron-specific network models of genetic regulatory programs in brain regions affected by LOAD. For these analyses, we utilized wholegenome gene expression and whole-genome genotyping data from two independent cohorts in the Accelerating Medicines Partnership - Alzheimer’s Disease (AMP-AD) consortium: the Mayo RNAseq Study (herein MAYO) and the Religious Orders Study and Memory and Aging Project (herein ROSMAP).

We first applied a deconvolution method to deconvolve bulk-tissue RNA sequencing (RNAseq) data from post-mortem brain regions and derive the neuron-specific gene expression signal. Although single-cell RNA sequencing (scRNAseq) studies have significantly advanced our understanding of cellular heterogeneity[23–25] and the discovery of novel cell populations[26, 27], as well as spurred developments of various computational analysis tools[28], network inference performance using scRNAseq data is still very poor. Due to the high volume of missing gene expression measures and the immaturity of current network methods dealing with these missing data, inferred network models using scRNAseq data yield a significant amount of uncertainty[29, 30], thus limiting the application of scRNAseq data in network inference. Alternatively, deconvolution of bulk-tissue RNAseq data has become increasingly popular in recent years as a complementary solution to the missing values in scRNAseq data[31–43], based on the core assumption that gene expression in bulk-tissue data is equal to the averaged gene expression of each cell type weighted by its relative population in the tissue. Deconvolution methods decompose bulk-tissue RNAseq data into gene expression of individual cell types by using celltype specific biomarker genes to implicitly estimate relative cell populations in the tissue. After deconvolution, the variances of the deconvoluted gene expression of each cell type become orthogonal to each other and can be analyzed independently[44].

To derive neuron-specific gene expression signals from the bulk-tissue RNAseq data from the MAYO and ROSMAP cohorts, we chose to employ the population-specific expression analysis (PSEA) method of deconvolution [44]. Whereas other popular deconvolution methods such as Cibersort [35], dtangle [45], DSA [31], or NNLS [46] can only estimate cell fraction in a bulk-tissue sample, the PSEA method directly estimates cell-type specific residuals from bulk-tissue RNAseq data. We demonstrated the robustness of the PSEA deconvolution method using random selection of neuronal biomarkers derived from scRNAseq studies [47–51].

After deconvolution, we applied a cutting-edge systems biology approach[52–54] to build causal network models of the neuronal component of AD by integrating the deconvoluted neuron-specific RNAseq data with the whole-genome genotype (SNP) data from the MAYO and ROSMAP datasets. We then agnostically identified neuron-specific gene regulatory network models and key genetic drivers (and thus potential therapeutic targets) predicted to modulate pathological Aβ and hyperphosphorylated tau accumulation in AD. To evaluate and ensure the robustness of our results, we performed the integrative analysis and key driver identification independently in the two cohorts and cross-validated the results at every step of the analysis. In total, we reconstructed 11 causal network models combined across the two separate analyses and predicted a total of 1,563 potential key drivers modulating neuronal network states and AD pathology under LOAD. To validate our network prediction, we prioritized 19 novel targets which replicated across our two cohorts for experimental validation. We used shRNA-mediated knockdown in human iPSC-derived neurons (iNs)[55–57] and measured levels of Aβ38, Aβ40, and Aβ42 as well as tau and p231-tau. Among the 19 novel targets, we identified 10 targets which affected Aβ (*JMJD6, NSF, NUDT2, DCAF12, RBM4, YWHAZ, NDRG4*, and *STXBP1*) and/or tau/p-tau levels (*JMJD6, FIBP*, and *ATP1B1*).

To further validate our network models and to provide insights into network connectivity, we measured the whole-genome RNA expression of iNs by RNAseq after knocking down each of the 19 targets and characterized differential gene expression (DE gene signatures) between each target shRNA and its associated controls. We validated network models by comparing the DE gene signature of each target to the downstream structure of the target in the networks, and we investigated pathways enriched by the gene knockdown DE signatures to shed light on the molecular mechanisms associated with LOAD, identifying our validated targets as upstream regulators of master regulators VGF and REST.

## Results

### An Integrative Systems Biology Approach for Constructing Single Cell-Type Regulatory Networks of AD

We developed an integrative network analysis pipeline to construct data-driven neuron-specific predictive networks of AD (Fig. 1). The overall strategy for elucidating the single cell-type gene network model depicted in Fig. 1 centers on the objective, data-driven construction of causal network models, which can be directly queried to identify the network components causally associated with AD as well as the master regulators (key drivers) of these AD-associated components. This model also predicts the impact of the key drivers on the biological processes and pathology involved in AD, moving us towards precision molecular models of disease. We previously developed this network reconstruction algorithm, i.e., predictive network, which statistically infers causal relationships between DNA variation, gene expression, protein expression, and clinical features measured in hundreds of individuals[21, 52, 58].

**Figure 1.**
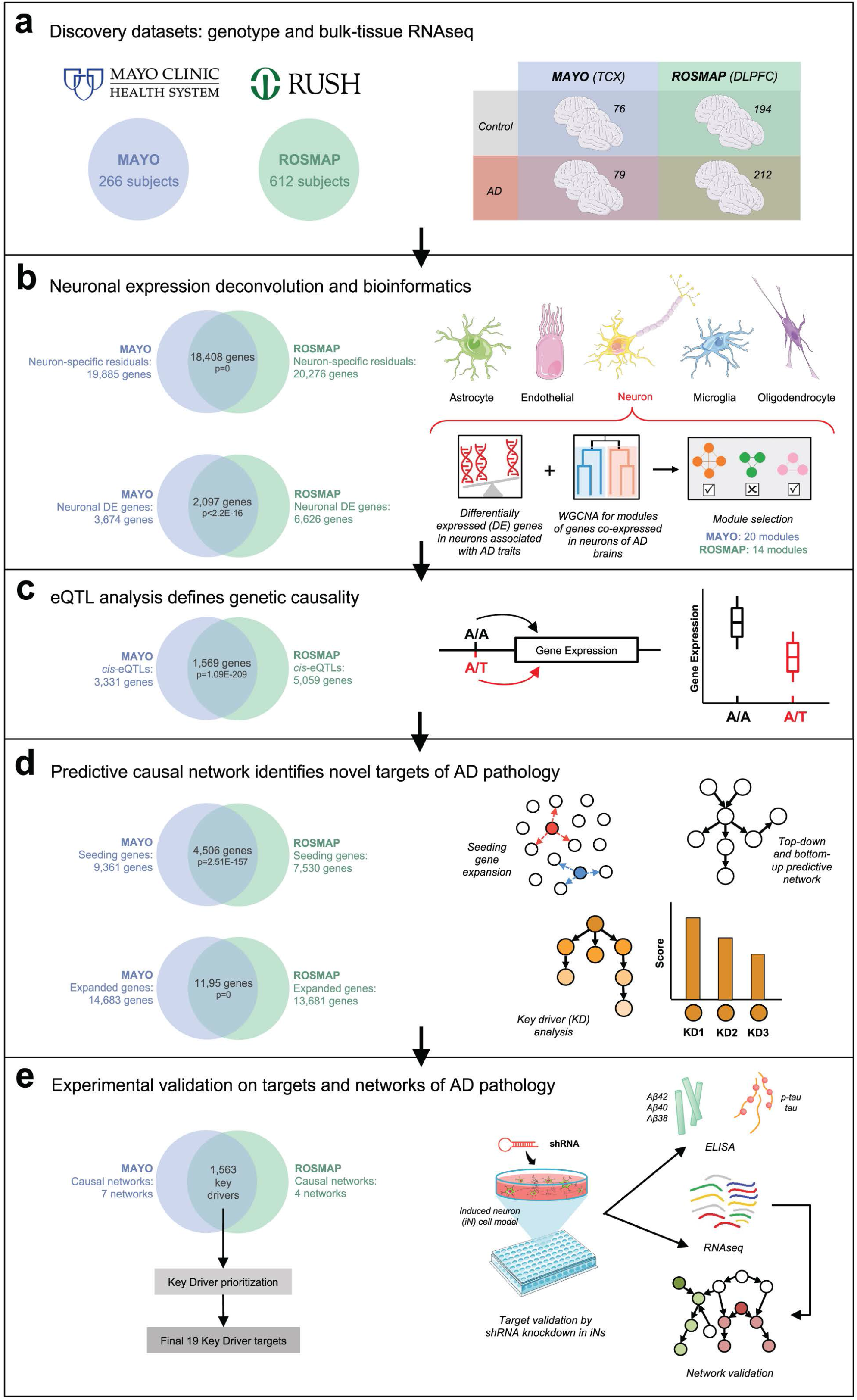
Integrative network analysis pipeline to construct data-driven neuron-specific predictive networks of AD and predict key drivers associated with AD pathology. **(a)** Discovery datasets include whole-genome genotype and RNAseq data of temporal cortex (TCX) from the MAYO cohort and dorsolateral prefrontal cortex (DLPFC) from the ROSMAP cohort in the NIH/NIA AMP-AD consortium. Numbers on the left indicate the total number of subjects in each dataset with quality-controlled matched genotype and RNAseq data used in this study, whereas numbers on the right (table) indicate the number of individuals of each phenotype used in the subsequent DE, co-expression module, and predictive network analyses. **(b)** Computational deconvolution of the bulk-tissue RNAseq data into 5 single cell-type RNAseq sets per cohort dataset, followed by differential expression analysis and weighted gene co-expression network analysis (WGCNA) in each cohort’s neuron-specific gene expression dataset. **(c)** mRNA expression and quantitative trait loci (expression-QTL, eQTL) association analysis in each dataset provides a source of systematic perturbation for network reconstructions. **(d)** Construction of neuron-specific predictive network models and identification of key drivers (master regulators) from each dataset. **(e)** Prioritization of key drivers targets from both datasets and experimental validation by shRNA-mediated gene knockdown in human iPSC-derived neurons. Venn diagrams at left on each panel indicate cross-validation at each step of the bioinformatics analyses performed independently in parallel for the MAYO and ROSMAP datasets, resulting in a single set of key driver targets. Statistical tests for each comparison are described in the text where relevant. Graphics not created by the authors were used with permission from Servier Medical Art (smart.servier.com).

The inputs required for our network analysis are the molecular and clinical data generated in the MAYO and ROSMAP populations, as well as first order relationships between these data such as quantitative trait loci (QTLs) associated with the molecular traits. These relationships are input as structure priors to the network construction algorithm as a source of perturbation, boosting the power to infer causal relationships at the network level, as we and others have previously shown [19, 21, 22, 58–66]. To focus on the component of AD that is intrinsically encoded in neurons, we identified the neuron-specific expression component in each cohort by applying the PSEA deconvolution algorithm[44] to the MAYO and ROSMAP transcriptomic datasets independently (Fig. S1, Step 1). We further focused on the molecular traits associated with AD by identifying differential gene expression (DE) signatures – comprised of several thousands of gene expression traits – between AD and cognitively normal (CN) samples for each dataset (Fig. S1, Step 2). To identify correlated gene expression traits associated with AD, we constructed gene co-expression networks for each dataset, and from these networks we identified highly interconnected sets of co-regulated genes (modules) that were significantly enriched for AD gene signatures (the significant DE genes) as well as for pathways previously implicated in AD (Fig. S1, Step 3). To obtain a final set of genes for input into the causal network construction process for each dataset, we combined genes in the co-expression network modules enriched for AD signatures (Fig. S1, Step 5-Module selection) and performed the pathFinder algorithm[58] to enrich the seeding gene set by including genes upstream and downstream of this set from a compiled pathway database (Fig. S1, Step 5-Seeding expansion). Note that we only include genes from the pathways without their interactions; the interactions among the final extended genes are solely inferred from each dataset.

With our input set of neuron-centered genes for the AD network constructions defined, we mapped expression-QTLs (eQTLs) for neuron-specific gene expression traits in each dataset to incorporate the eQTLs as structure priors in the network reconstructions, given that they provide a systematic perturbation source that can boost the power to infer causal relationships (Fig. S1, Step 4)[22, 59, 60, 62–67]. The input gene set and eQTL data were then processed by an ensemble of causal network inference methods including Bayesian networks and our recently developed top-down and bottom-up predictive networks [52, 58, 68, 69], in order to construct probabilistic causal network models of AD independently in the MAYO and ROSMAP cohorts (Fig. S1, Step 6). We next applied a statistical algorithm to detect key driver genes in each given network structure[70] and to identify and prioritize master regulators in the AD networks (Fig. S1, Step 7). These key drivers derived from the individual networks across datasets were then pooled and prioritized based on ranking scores of impact and robustness (Methods), resulting in a final group of 19 top-prioritized key drivers for which we performed functional validation in a human induced pluripotent stem cell (iPSC) derived neuron system. The entire analysis workflow for the independent datasets, resulting in this final group of replicated targets, is illustrated in Fig. 1.

### The Mayo Clinic and ROSMAP Study Populations and Data Processing

Our causal network pipeline starts by integrating whole-genome genotyping and RNAseq data generated from patients spanning the complete spectrum of clinical and neuropathological traits in AD. We used patient data from two separate cohorts within the AMP-AD consortium: temporal cortex (TCX) data from 266 subjects in MAYO[71–73] and dorsolateral prefrontal cortex (DLPFC) data from 612 subjects in ROSMAP[20, 74–76] (Fig. 1a). We processed matched genotype and RNAseq data separately in each dataset (Fig. 1, Fig. S1; Methods).

CNS tissue consists of various cell types, including neurons, glia, and endothelial cells. To discover key network drivers specific to a single cell type in the CNS and study their contribution to AD in that specific cell type, we utilized verified single-cell marker genes to directly deconvolve bulk-tissue gene expression data into cell-type specific gene expression for the five major cell types in the CNS: neurons, microglia, astrocytes, endothelial cells, and oligodendrocytes (Methods). In this study, we focused on investigating the role of neuronal cells in AD, as they are the primary cell type affected by AD pathogenesis [77–79]. After normalizing the bulk-tissue RNAseq data, we performed variance partition analysis (VPA)[80] to evaluate the contributions of cell-type specific markers as well as demographic, clinical, and technical covariates (such as batch effects) to the gene expression variance before performing any covariate adjustment (Fig. S2a,b). The cell-type specific marker genes used for neurons, microglia, astrocytes, endothelial cells, and oligodendrocytes were *ENO2, CD68, GFAP, CD34*, and *OLIG2*, respectively, as previously published [72]. The VPA results reflect the prominent effect of CNS cell types on the variance of the brain RNAseq data. In the MAYO dataset, the additional covariates used in the VPA included exonic mapping rate, RNA integrity number (RIN), sequencing batch, diagnosis, age at death, tissue source, *APOE* genotype, and sex. In the ROSMAP dataset, we were able to include the same covariates with the exception of tissue source and the addition of age at first AD diagnosis, post-mortem interval (PMI), education, and study (ROS or MAP).

We then performed covariate adjustment and deconvolution using the PSEA method[44] in each dataset, calculating gene expression residuals using a linear regression model to adjust the normalized bulk-tissue expression data with demographical and technical covariates as well as the cell-type specific markers. Cell-type specific gene expression, including the neuron-specific component, was directly derived by adding the estimated variance of each cell type to the residual (Methods), avoiding the need to first estimate the cell population from bulk tissue data, which could induce approximation errors. We then repeated VPA in the neuron-specific residuals of each dataset to demonstrate that our deconvolution and covariate adjustment methods properly capture the neuronal component while removing potential confounds such as batch effect, age, and sex (Fig. S2c,d). Finally, to justify the use of single cell-type specific markers for deconvolution by the PSEA method, we performed a set of analyses comparing multiple cell-type specific biomarker lists (derived from existing scRNAseq studies) to each other, to our AD residuals, and to the AMP-AD Agora list of potential therapeutic targets in AD (Fig. S3; Methods), as well as a robustness analysis demonstrating that our neuron-specific residual derived from *ENO2* expression represents a robust neuronal component in the bulk-tissue RNAseq data when compared to random selections of multi-gene neuronal biomarkers derived from these scRNAseq datasets in AD (Fig. S4; Methods).

### Identifying AD-Associated Gene Signatures in Neurons and Mapping Their eQTLs

To identify an AD-centered set of neuronal gene expression traits, we performed differential expression (DE) analysis using the deconvoluted neuron-specific expression residuals in the MAYO and ROSMAP cohorts (Methods). In comparing expression data between AD and cognitively normal (CN) controls (MAYO-TCX: 79 AD, 76 CN; ROSMAP-DLPFC: 212 AD, 194 CN), there were 3,674 significant DE neuron-specific genes in the MAYO dataset (hereby “MAYO-neuron”) and 6,626 neuron-specific DE genes in the ROSMAP dataset (hereby “ROSMAP-neuron”) (Fig. 2a,b, FDR<0.05; Fig. S5). There were 2,097 significant DE genes overlapping between the two datasets (Supplementary Table S1, Fisher Exact Test, odd ratio=3.9784, p-value<2.2E-16), thus cross-validating the neuron-specific DE signatures independently derived from the two cohorts.

**Figure 2.**
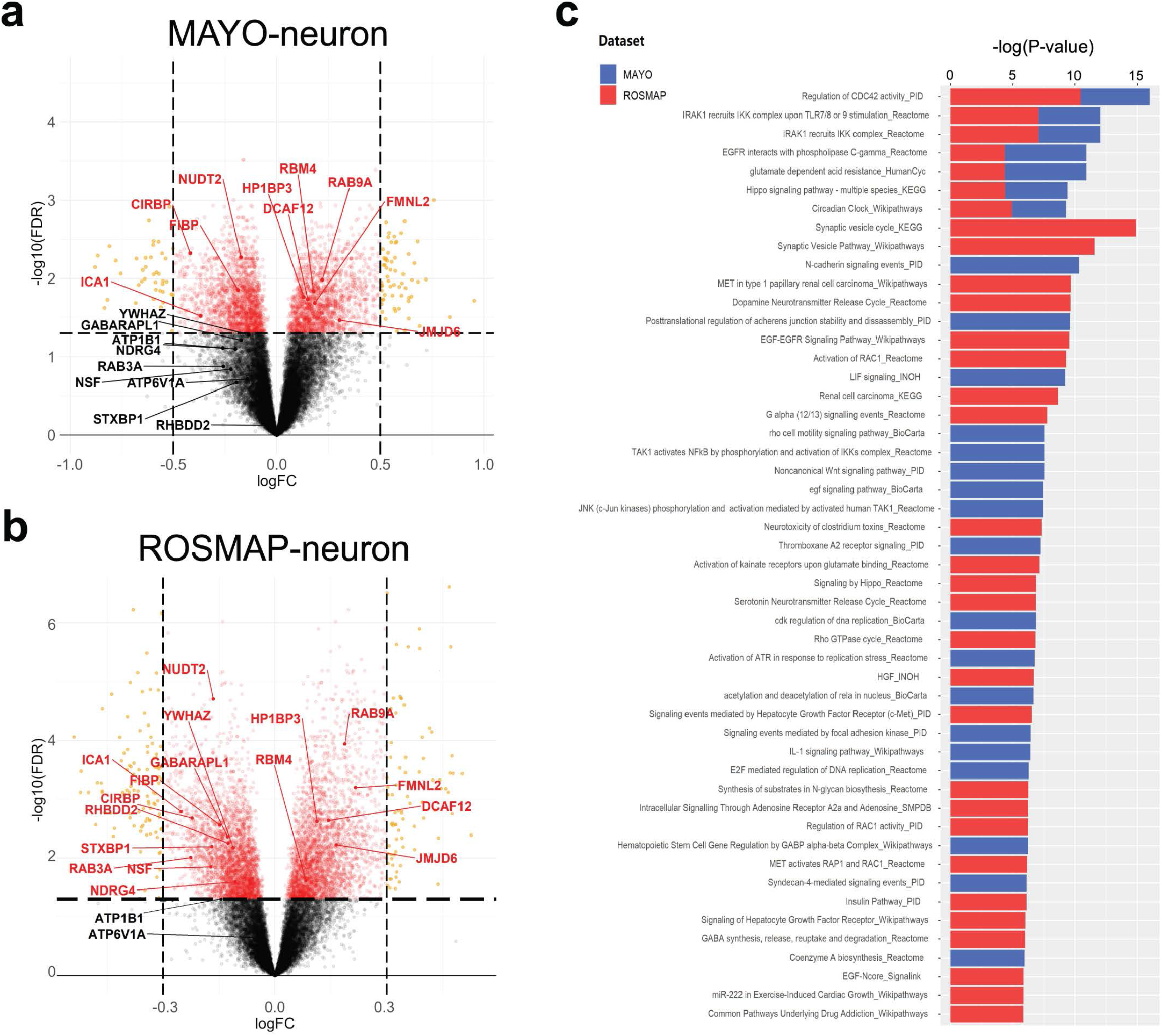
Neuron-specific gene expression signatures in AD. Differential expression (DE) analysis of deconvoluted neuron-specific residuals identifies a robust DE signature associated with the difference between AD patients and cognitively normal controls. Volcano plots **(a,b)** show all significantly up- and down-regulated genes with cutoffs on absolute log fold-change greater or less than 0.5 for MAYO-neuron (**a**) and 0.3 for ROSMAP-neuron (**b**). Significance was assessed by t-test with FDR<0.05. Gene symbols are highlighted for the 19 key drivers prioritized for experimental validation *in vitro*. **(c)** Pathway enrichment analysis with human ConsensusPathDB (CPDB) on the neuron-specific DE expression signatures reveals dysregulated biological processes associated with AD. Significance was assessed by Fisher’s exact test with p-value<0.05. Detailed statistical results of DE genes and enriched pathways are summarized in Supplementary Tables S1 and S2, respectively.

To examine the biological processes that are dysregulated in AD cases versus controls as reflected in the DE signatures, we performed pathway enrichment analysis on the MAYO-neuron and ROSMAP-neuron gene sets using Human ConsensusPathDB (CPDB)[81–85]. We identified 75 and 73 enriched pathways in each dataset, respectively, with 7 pathways significantly dysregulated in both datasets (Fig. 2c, Supplementary Table S2, p-value<0.05). These signatures were enriched for a number of cellular/molecular pathways, including those involving CDC42[86–88], IRAK/IKK[89–91], EGFR/PLCG[92], GAD[93], Hippo[94], and clock genes[95, 96], some of which have been implicated and/or interrogated in AD previously. Additional pathways of note implicated by a single cohort (MAYO or ROSMAP) with known relevance to amyloid and/or tau pathology include those related to NF-κB activation[97, 98] and N-cadherin signaling[99, 100].

We further validated our neuron-specific DE signatures in AD, which were derived from deconvoluted bulk-tissue RNAseq data by comparing our MAYO-neuron and ROSMAP-neuron DE genes with the excitatory and inhibitory neuronal signatures identified by a separate study which generated scRNAseq data from the same ROSMAP cohort[101]. We employed the sampling-based method described in this study[101] and found highly significant enrichment between our MAYO-neuron and ROSMAP-neuron DE signatures and their excitatory neuron signature (p=5.69E-11 and p=2.97E-18, respectively) as well as their inhibitory neuron signature (0.019 and 0.0025, respectively), demonstrating significant correlation between our deconvoluted neuron-specific DE signatures and scRNAseq-derived neuronal signatures in AD[101] (Fig. S4). The greater excitatory-neuronal enrichment among our deconvoluted neuron-specific DE signatures is consistent with the single-cell transcriptomics study in AD[101] and similarly suggests that our deconvoluted RNAseq datasets capture the aberrant increases in neuronal excitotoxicity associated with AD in humans[102].

Another critical input for the construction of Bayesian network and causal predictive network models are the eQTLs, leveraged as a systematic source of perturbation to enhance causal inference among molecular traits. This is an approach we and others have demonstrated across a broad range of diseases and data types[22, 59–63, 65–67, 69, 103–116]. We mapped *cis*-eQTLs by examining the association of neuron-specific expression traits with genome-wide genotypes [18, 117–120] assayed in the MAYO and ROSMAP cohorts (Methods). In the MAYO- and ROSMAP-neuron sets, 3,331 (16.8%) and 5,059 (25.0%), respectively, of the residual genes were significantly correlated with allele dosage (FDR<0.01) (Supplementary Table S3). Of the *cis*-eQTLs detected in each cohort, 1,569 genes were overlapping between the two sets (47% of MAYO *cis*-eQTLs and 31% of ROSMAP *cis*-eQTLs, Fisher’s Exact Test, p-value=3.31E-242), providing further validation of the two independent cohorts.

### Neuronal Co-expression Networks Associated with LOAD

While DE analysis can reveal patterns of neuron-specific expression associated with AD, the power of such analysis to detect a small-to-moderate expression difference is low. To complement the DE analyses in identifying the input gene set for the causal network, we clustered the neuronal gene expression traits into data-driven, coherent biological pathways by constructing coexpression networks, which have enhanced power to identify co-regulated sets of genes (modules) that are likely to be involved in common biological processes under LOAD. We constructed co-expression networks on the AD patients within each dataset after filtering out lowly expressed genes (Methods), resulting in the MAYO-neuron co-expression network consisting of 20 modules ranging in size from 30 to 6,929 gene members and the ROSMAP-neuron coexpression network consisting of 14 modules ranging from 34 to 6,604 gene members (Fig. 3a).

**Figure 3.**
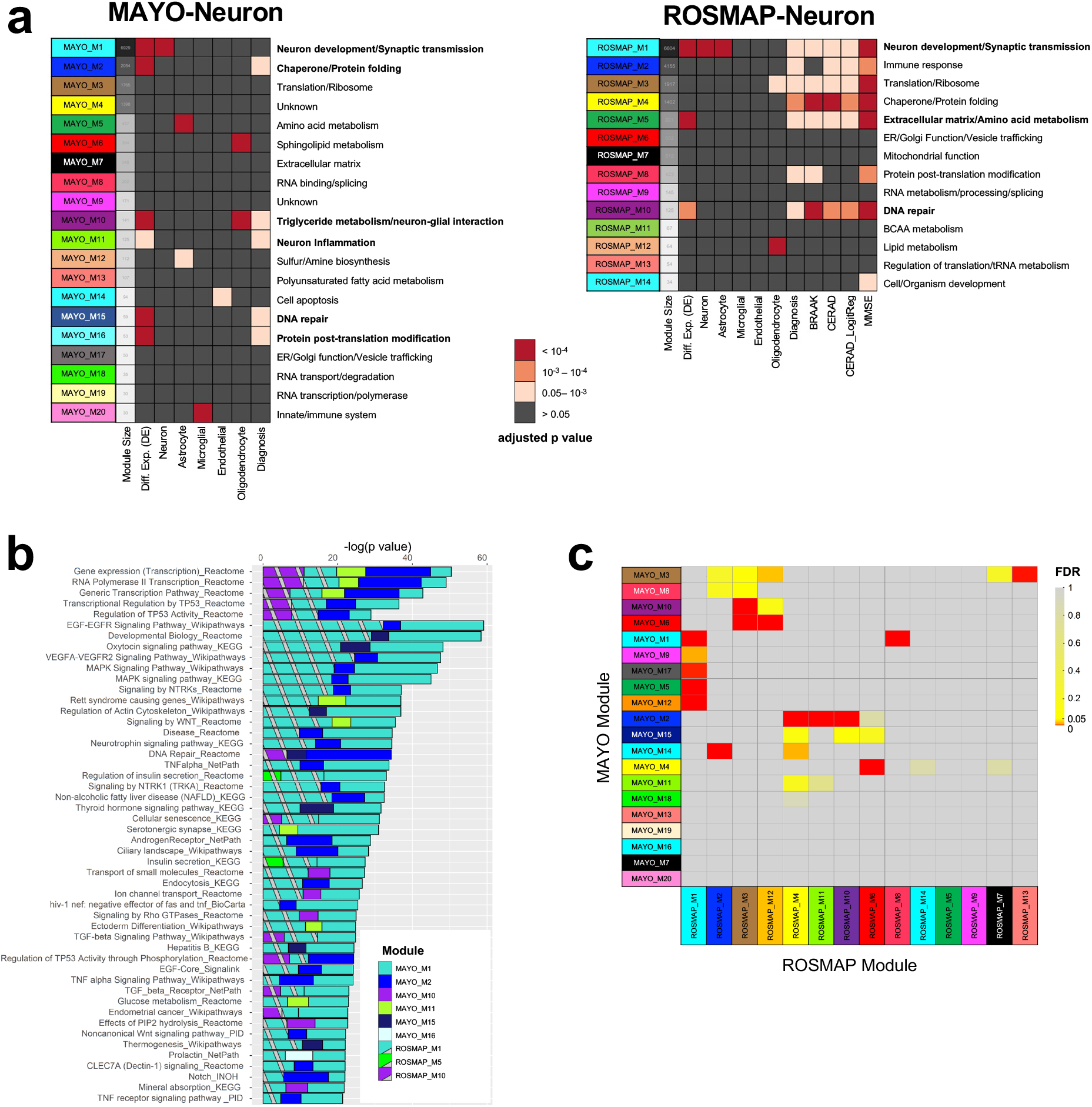
Neuron-specific co-expression analysis identifies robust gene modules enriched for biological processes associated with AD. **(a)** Neuron-specific co-expression network analysis in the MAYO and ROSMAP cohorts identifies gene modules associated with AD in each dataset. Module functions for each dataset are characterized by significantly enriched biological processes (Supplementary Table S4), with bold text indicating neuron-specific modules selected for further analysis. Each module was evaluated based on enrichment for neuron-specific DE genes, for scRNAseq derived neuron-specific biomarker genes, and for categories of available AD traits (MAYO and ROSMAP diagnosis by ANOVA; BRAAK, CERAD, and MMSE by linear regression). We also evaluated enrichment for scRNAseq-derived biomarker genes for other four major CNS cell types (microglia, astrocytes, endothelial cells, and oligodendrocytes) to cross-validate that modules enriched for neuron-DE genes were not enriched for other cell types. Significance was assessed by Fisher’s exact test with adjusted p-value<0.05. (**b**) Pathway enrichment analysis identified robust pathways replicated in both cohorts associated with AD among the selected neuron-specific modules in the two datasets. (**c**) Cross-validation of neuron-specific co-expression modules between MAYO and ROSMAP identifies pairs of modules with significantly overlapping gene members (FDR<0.05).

To evaluate the functional relevance of the each cohort’s neuron-specific modules to AD pathology, we performed enrichment analysis of each module for its AD-associated neuronal DE signatures, known single-cell marker genes for the 5 major cell types in the CNS[50], and categories of AD traits available from its respective cohort (Fig. 3a). From these enrichment results, we identified neuron-specific modules associated with AD DE genes (FDR<0.05) from the two co-expression networks: M1, M2, M10, M11, M15, and M16 from MAYO and M1, M5, and M10 from ROSMAP.

To further characterize the biological processes involved in the co-expression modules from each dataset, we performed pathway enrichment analysis to identify overrepresented biological processes within and across the modules (Fig. 3b, Supplementary Table S4). Out of the selected AD-associated modules from the MAYO- and ROSMAP-neuron co-expression networks, respectively, there were 36 and 16 significantly enriched pathways (FDR<0.05) based on the Human ConsensusPathDB (CPDB) database, with 11 enriched pathways overlapping between the two datasets (Fig. 3b, Fisher’s Exact Test, OR=383.87, p-value<2.2E-16). In comparing all pairs of modules between the datasets, we identified 17 module pairs with significant overlap of gene members (Fig. 3c, FDR<0.05 for Fisher’s Exact Test), demonstrating the robustness of the two independent co-expression networks.

### Ensemble of Neuronal Causal Networks of Genetic Regulations Identifies Pathological Pathways and Key Drivers for Neuronal Function in AD

The ultimate goal of this study was to identify upstream master regulators (key drivers) of neuronal pathways that contribute to AD. Following our DE, eQTL, and co-expression network analyses, we built an ensemble of causal network models – including standard Bayesian networks[20, 22] and state-of-the-art predictive network models[19, 21, 58] – by integrating the eQTLs and deconvoluted neuron-specific RNAseq residuals.

We first pooled all genes from the selected AD-associated modules per dataset (six MAYO-neuron modules and three ROSMAP-neuron modules, indicated in Fig. 3a) to create a seeding set of genes for each cohort for input into the network models. This resulted in 9,361 seeding genes from the MAYO-neuron co-expression network and 7,530 seeding genes from the ROSMAP-neuron co-expression network. We note an overlap of 4,506 genes between the two seeding gene sets (48.1% of MAYO and 59.8% of ROSMAP, Fisher’s Exact Test, p-value<2.2e-16), indicating the reproducibility of these analyses across the two independent datasets. To further improve the robustness for our network models, we also expanded each set of seeding genes by including their known upstream and downstream genes in each cohort’s co-expression network, extracted from signaling pathway databases using the pathFinder algorithm[58] (Methods; note that we did not include the gene-gene interactions as prior edge information for network construction). Co-expression network modules are only sensitive to linear relationships between pairs of genes, whereas non-linear gene regulations will not be captured by coexpression analysis. This expansion step thus includes genes in the same pathways as the seeding genes which otherwise failed to be included in the same module derived from the coexpression networks, resulting in 14,683 expanded genes from MAYO-neuron, 13,681 expanded genes from ROSMAP-neuron, and an overlap of 11,952 genes between the two expanded gene sets. The use of both the seeding gene set and the expanded gene set for analysis of the MAYO and ROSMAP datasets therefore increases the power to build robust networks and to discover high-confidence neuronal key drivers associated with AD pathology.

We also incorporated *cis*-eQTL genes into each network as structural priors. As *cis*-eQTLs causally affect the expression levels of neighboring genes, they can serve as a source of systematic perturbation to infer causal relationships among genes[21, 52, 58, 67]. Of the 3,331 and 5,059 unique *cis*-eQTL genes identified in the MAYO- and ROSMAP-neuron datasets, respectively, 687 and 1978 overlapped with the seeding gene set and 2,162 and 2,998 overlapped with the expanded gene set. We finally proceeded to build Bayesian networks and predictive networks using the two sets of genes per dataset – i.e., 9,361 seeding and 14,683 expanded genes for the MAYO dataset and 7,530 seeding and 13,681 expanded genes for the ROSMAP dataset – and incorporating each dataset’s *cis*-eQTL genes as structural priors.

Since structure learning is a heuristic and stochastic process, we applied a wide range of cut-offs on the posterior probability of edges to derive sets of robust Bayesian and predictive network structures for each dataset. For the MAYO-neuron seeding gene set, we built Bayesian networks and applied two posterior probability cut-offs (0.4/0.5, Methods) to get two MAYO-neuron Bayesian networks (MAYO-Neuron-BayesNet-Seed-1/-2) which were comprised of 9,111/9,044 genes, respectively. In addition, we built predictive networks with the same two posterior probability cut-offs (0.4/0.5) to derive two MAYO-neuron predictive networks (MAYO-Neuron-PredNet-Seed-1/-2), which also included 9,111/9,044 genes, respectively. For the MAYO-neuron expanded gene set, we built predictive networks and chose three posterior probability cut-offs (0.5/0.6/0.7) to get three MAYO-neuron predictive network models (MAYO-Neuron-PredNet-Expanded-1/-2/-3), which were comprised of 14,238/13,926/13,365 genes, respectively. For the ROSMAP-neuron seeding gene set, we built Bayesian networks and applied two cut-offs (0.3/0.4) to derive two Bayesian networks (ROSMAP-Neuron-BayesNet-Seed-1/-2) which consisted of 6,786/6,756 genes, respectively. For the ROSMAP-neuron expanded gene set, we built two predictive networks and chose two cut-offs (0.3/0.4) to build two predictive networks (ROSMAP-Neuron-PredNet-Expanded-1/-2) consisting of 12,147/12,074 genes, respectively. Thus, in total from the MAYO and ROSMAP datasets, we derived 11 networks for the inference of a robust set of key drivers, using several different network reconstruction methods, network gene sets, and posterior cut-offs. We demonstrate 2 of the final 11 causal network models in Fig. 4a,b (MAYO-Neuron-PredNet-Expanded-1 and ROSMAP-Neuron-PredNet-Expanded-1), and the remaining 9 causal networks are shown in Fig. S6.

**Figure 4.**
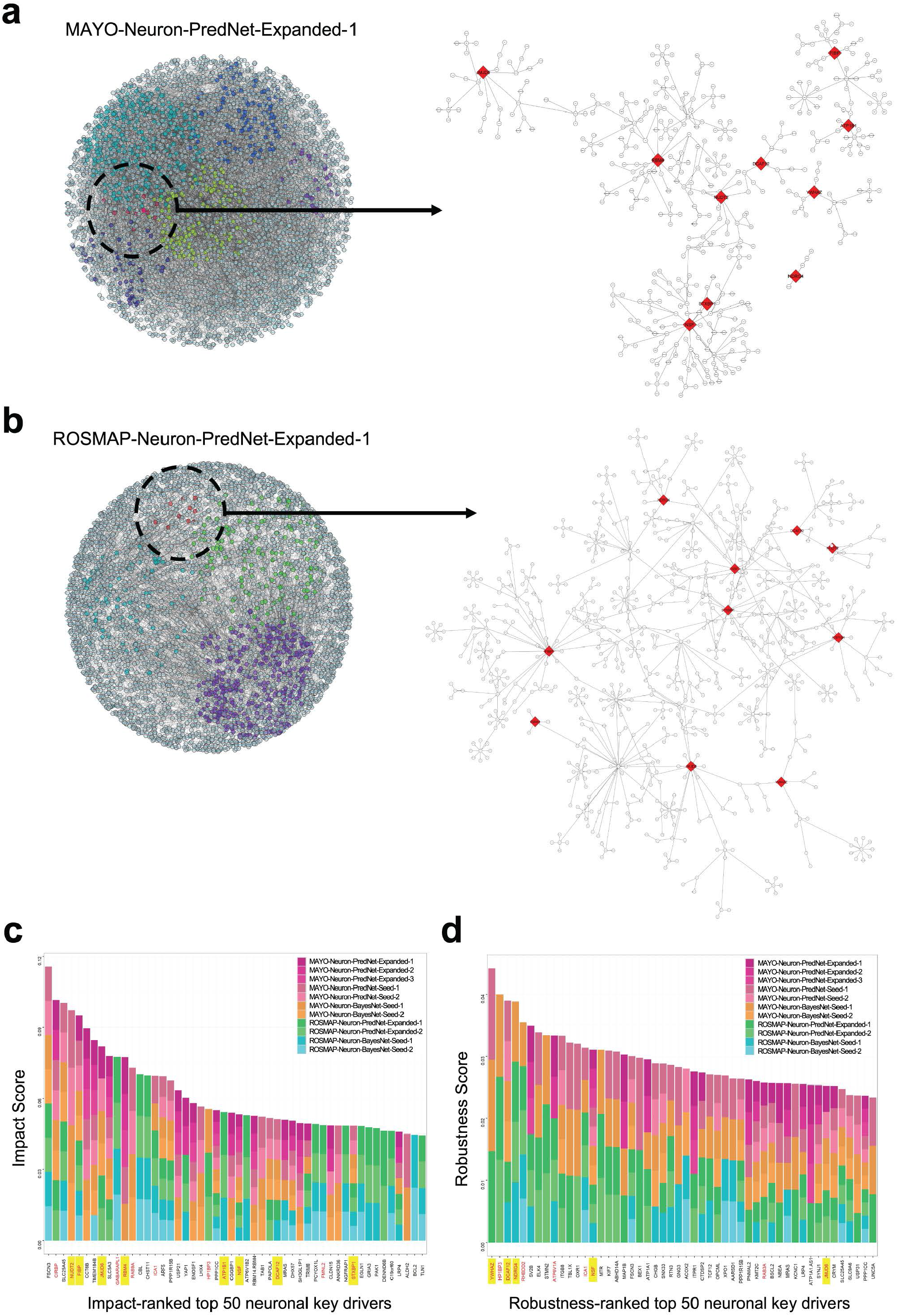
Neuron-specific causal network analyses identify molecular mechanisms and key driver targets associated with AD. (**a,b**) Two predictive networks out of the final 11 neuron-specific causal Bayesian and predictive network models derived from the MAYO **(a)** and ROSMAP **(b)** seeding and expanded gene sets. The MAYO and ROSMAP networks shown here were built from their respective expanded gene sets with posterior probability cut-offs of 0.5 and 0.3, respectively. The 10 key driver targets which were validated *in vitro* are highlighted (red), along with their neighboring downstream subnetworks. **(c,d)** The top 50 out of 1,563 total key driver targets ranked individually according to impact **(c)** and robustness **(d)** scores across the 11 independent MAYO-neuron and ROSMAP-neuron Bayesian and predictive networks. Red text indicates prioritized key drivers; yellow highlights those which were validated *in vitro*.

### Identification and Prioritization of Neuronal Key Drivers Regulating AD Pathology

Having generated the causal predictive networks from the MAYO-neuron and ROSMAP-neuron datasets, we applied Key Driver Analysis (KDA)[70] to derive a list of key driver genes from each network. KDA seeks to identify genes in a causal network which modulate network states; in the present analysis, we applied KDA to identify genes causally modulating the network states of our neuron-specific Bayesian and predictive network models. In total, we identified 1,563 key driver genes across the 11 independent networks.

To prioritize key drivers for further investigation, we first ranked the 1,563 initial key driver targets according to two separate measures: an impact score and a robustness score (Methods). Briefly, the impact score is a predicted value quantifying the regulatory impact of a given key driver on its downstream effector genes associated with AD pathology. Intuitively, the shorter a path from a key driver to its downstream effectors in a network – with less other parental co-regulators along the same path – the greater the impact of this target on its effectors in that network. Robustness score is reflective of the number of datasets (MAYO and/or ROSMAP), gene sets (seeding and/or expanded), and network models (Bayesian and/or predictive) by which a key driver is replicated. After ranking the total 1,563 neuron key drivers according to each score, we focused on the top 50 key drivers in each ranked list (Fig. 4c,d).

We then performed a series of steps to prioritize a final group of key driver targets for *in vitro* experimentation out of the ensemble of the top 50 ranked candidates for each score. We first calculated the replication frequency across the two ranked lists and identified 11 replicated targets, indicating robustness across these two independent ranking scores, and 39 unique targets in each ranked list (78 total). For the 11 replicated targets, we removed any which ranked lower than 15 in both scores, resulting in 7 top-ranked targets (*ICA1, NSF, FSCN3, HP1BP3, DCAF12, JMJD6*, and *SLC25A45*) which were replicated in both lists and ranked within the top 15 in one or both scores. Next, for the remaining 78 unique targets, we first selected the top 3 unique targets from each ranked list (*CIRBP, NUDT2*, and *FIBP* for impact score; *YWHAZ, NDRG4*, and *RHBDD2* for robustness score). To further select targets from the remaining 36 neuron-specific targets in each ranked list (72 total), we identified 4 targets (*GABARAPL1, ATP1B1, ATP6V1A*, and *RAB3A*) which were previously nominated to the AMP-AD Agora portal list based on separate data-driven network analysis using the bulk-tissue RNA-seq data in the MAYO and ROSMAP datasets with the same approach as this study[121]. Finally, to balance our selection strategy, we selected an additional 4 targets (*RBM4, RAB9A, FMNL2*, and *STXBP1*) out of the lower-to-middle ranked top 50 unique targets based on the availability of proper constructs.

In summary, we prioritized a group of 19 targets for experimental validation *in vitro* (highlighted red in Fig. 4c,d) by selecting the top-ranked replicated targets across the two scores (we note that *SLC25A45* and *FSCN3* were excluded at this stage due to lack of proper constructs), 6 top-ranked unique targets (top 3 from each score), 4 targets overlapping with prior data-driven nominations to the AMP-AD Agora list, and 4 lower-to-middle ranked targets.

### Validation of AD-Associated Function of Neuronal Key Drivers by Knockdown in Human Neurons

We next aimed to test the functional consequences of perturbation of the top candidate driver genes in human neurons. Healthy control human induced pluripotent stem cells (iPSCs) were differentiated to a neuronal fate using the well-established NGN2 direct differentiation protocol[56], which rapidly generates highly homogenous populations of induced neurons (iNs) which are cortical layer 2/3-like glutamatergic neurons[56, 57, 122]. By two weeks in culture, iNs are post-mitotic, electrically active, and express a full array of synaptic markers [56, 122]. In order to perturb the expression of the top 19 candidate key driver genes, we obtained sets of validated short hairpin RNA (shRNA) constructs packaged in lentivirus, with each set containing three constructs against each selected gene (Broad Institute). At day 17 of differentiation, iNs were transduced with lentivirus encoding a single shRNA, alongside control cells which either received empty virus or were not transduced (fresh media only). Media were exchanged on all cells 18 hours later. Five days following transduction (day 22 of differentiation), conditioned media were collected, and cells lysed, either to collect RNA for RNAseq or to harvest protein for analyses of Aβ and p-tau/tau, similar to our previous study of LOAD GWAS hits[123]. All Aβ and tau data were normalized to total protein in the cell lysate per well, and all data for each shRNA knockdown were additionally normalized to the average of control conditions (empty vector and no transduction) (Fig. 5).

**Figure 5.**
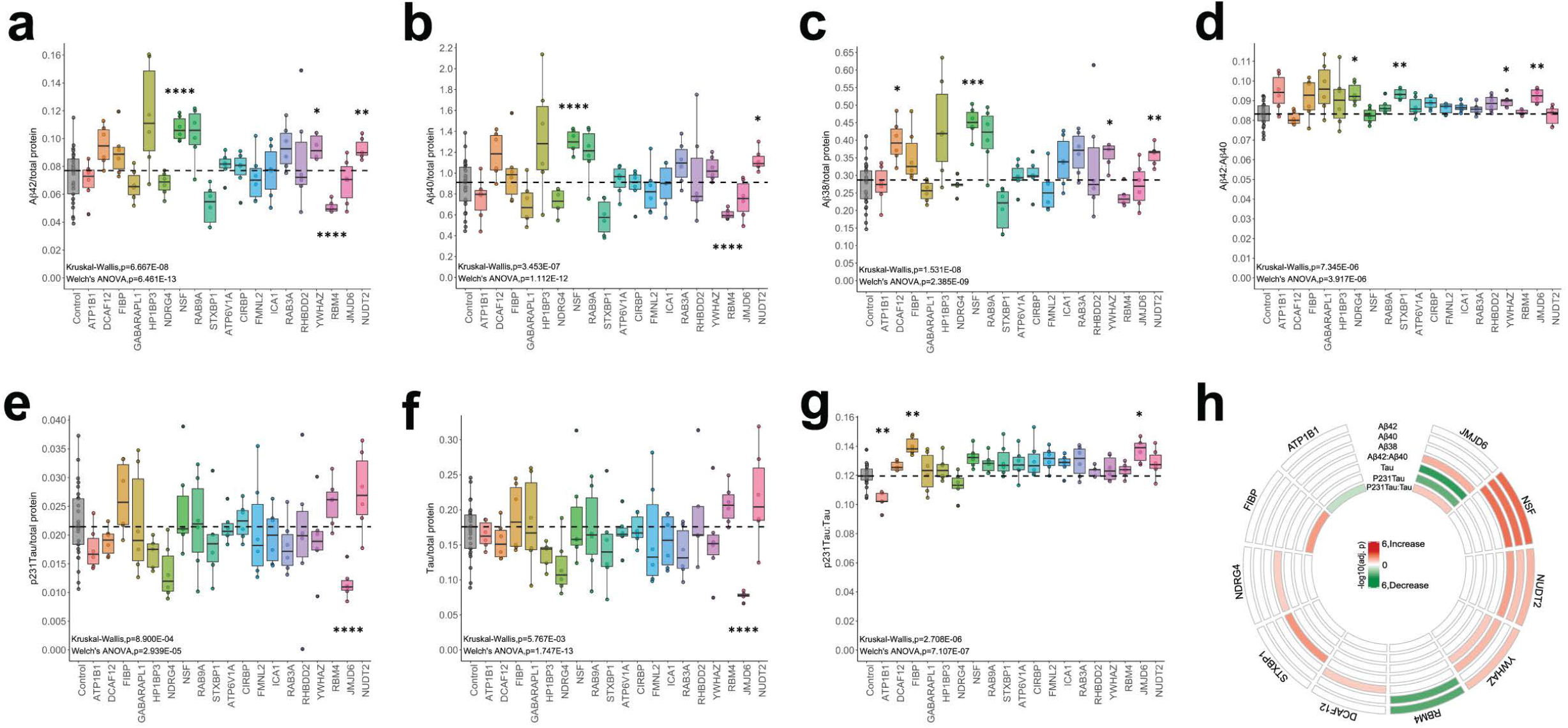
Human iPSC-derived neurons show altered Aβ species and tau/phospho-tau levels following shRNA-mediated knockdown of selected target genes. **(a-c)** Secretion of Aβ42 (**a**), Aβ40 (**b**), and Aβ38 (**c**) was measured in the conditioned media by ELISA (MSD), normalized to the average of controls (no transduction and empty vector) as well as to total protein in the neuronal cell lysate. The ratio of Aβ40:42 was also calculated **(d)**. **(e-f)** p231-tau (**e**) and total tau (**f**) were measured in cell lysates by ELISA (MSD), normalized to the average of controls (no transduction and empty vector) as well as to total protein in the cell lysate. The ratio of p231-tau to total tau was also calculated (**g**). For all panels, the black dashed line indicates the median for control conditions, and the black bar in each boxplot denotes the median of each gene knockdown condition. For each target gene, 3 shRNA constructs were used; each dot represents data from one well. A two-step statistical test was employed: first, we performed Welch’s ANOVA with unequal variance to detect significant differences across conditions for each measured parameter, with an additional non-parametric Kruskal-Wallis ANOVA to confirm the significance. Second, Dunnett’s T3 test (in Prism 9.0) with multiple testing correction was used to compare each target shRNA to the control condition for each parameter. *adj-p<0.05, **0.001<adj-p<0.05, ***0.0001<adj-p<0.001, ****adj-p<0.0001. **(h)** Circus plot summarizing the effects of the 10 key driver targets found to modulate Aβ42, 40, 38, Aβ42:40, tau, p231-tau and p231-tau:tau. Significance was assessed by -log10(Dunnett’s T3 adjusted p-value); red indicates that knockdown of the target significantly increased the given measurement value, whereas green indicates that knockdown significantly decreased the value. Detailed results and statistics are summarized in Supplementary Table S5.

Aβ38, 40, and 42 levels were measured in conditioned media from the transduced and control iNs using the Meso Scale Discovery (MSD) Triplex ELISA platform. Of the 19 genes tested, knockdown of 11 genes had no significant effect on the levels of any Aβ peptides measured nor the ratio of Aβ42 to Aβ40 (Fig. 5a-d). However, targeted knockdown of *YWHAZ* significantly raised Aβ42 peptide levels, knockdown of *DCAF12* and *YWHAZ* increased Aβ38 levels, and knockdown of *NSF* and *NUDT2* significantly increased levels of all three Aβ peptides measured (Aβ38, 40, and 42) (Fig. 5a-c; Dunnett’s T3 adjusted p-value<0.05). On the other hand, knockdown of *RBM4* significantly reduced levels of both Aβ42 and Aβ40 (Fig. 5a,b; Dunnett’s T3 adjusted p-value<0.05). Lastly, knockdown of *NDRG4, STXBP1, YWHAZ*, and *JMJD6* resulted in a significant elevation of the putatively neurotoxic Aβ42 to 40 ratio [124, 125](Fig. 5d; Dunnett’s T3 adjusted p-value<0.05).

We also examined levels of tau species in the transduced and control iN lysates using an MSD ELISA measuring both total tau and phospho-tau (Thr231). Knockdown of 16 of the 19 candidate genes tested had no significant effect on the levels of tau, p231-tau, or the neurotoxic ratio of p231-tau to tau (Fig. 5e-g). However, targeted knockdown of *JMJD6* significantly decreased the levels of both p231-tau and tau (Fig. 5e,f; Dunnett’s T3 adjusted p-value<0.05). We also note that knockdown of *NSF* approached significance of increased levels of p231-tau (Fig. 5e; Dunnett’s T3 adjusted p-value=0.075). Finally, knockdown of *FIBP* and *JMJD6* resulted in significant elevation of the p231-tau to tau ratio, while knockdown of *ATP1B1* significantly lowered this ratio (Fig. 5g; Dunnett’s T3 adjusted p-value<0.05).

In summary, here we confirm modulation of AD endophenotypes in human iNs following independent reduction of the expression of 10 different genes out of the top 19 predicted key driver targets (Fig. 5h). Detailed statistical results are included in Supplementary Table S5.

### Validation of AD-Associated Networks and Pathways by RNAseq of Human Neurons Following Targeted Gene Knockdown

To validate the network structure, we repeated shRNA-mediated knockdown of each of the 19 target key drivers in another set of cultured control iNs and subsequently measured gene expression by RNAseq. For each of the 10 AD endophenotype modulating targets, we derived a differential expression (DE) signature from the RNAseq data (Fig. 6a-j, Supplementary Table S6). Next, we extracted the downstream (sub)network of each of those 10 targets from the MAYO- and ROSMAP-neuron networks and evaluated the enrichment of the knockdown DE signature by the downstream subnetworks for each target. We found that 8 out of the 10 DE signatures were enriched by the downstream subnetworks of their corresponding target (Fig. 6k), validating that our network models capture a significant portion of molecular processes and pathways at the neuron level.

**Figure 6.**
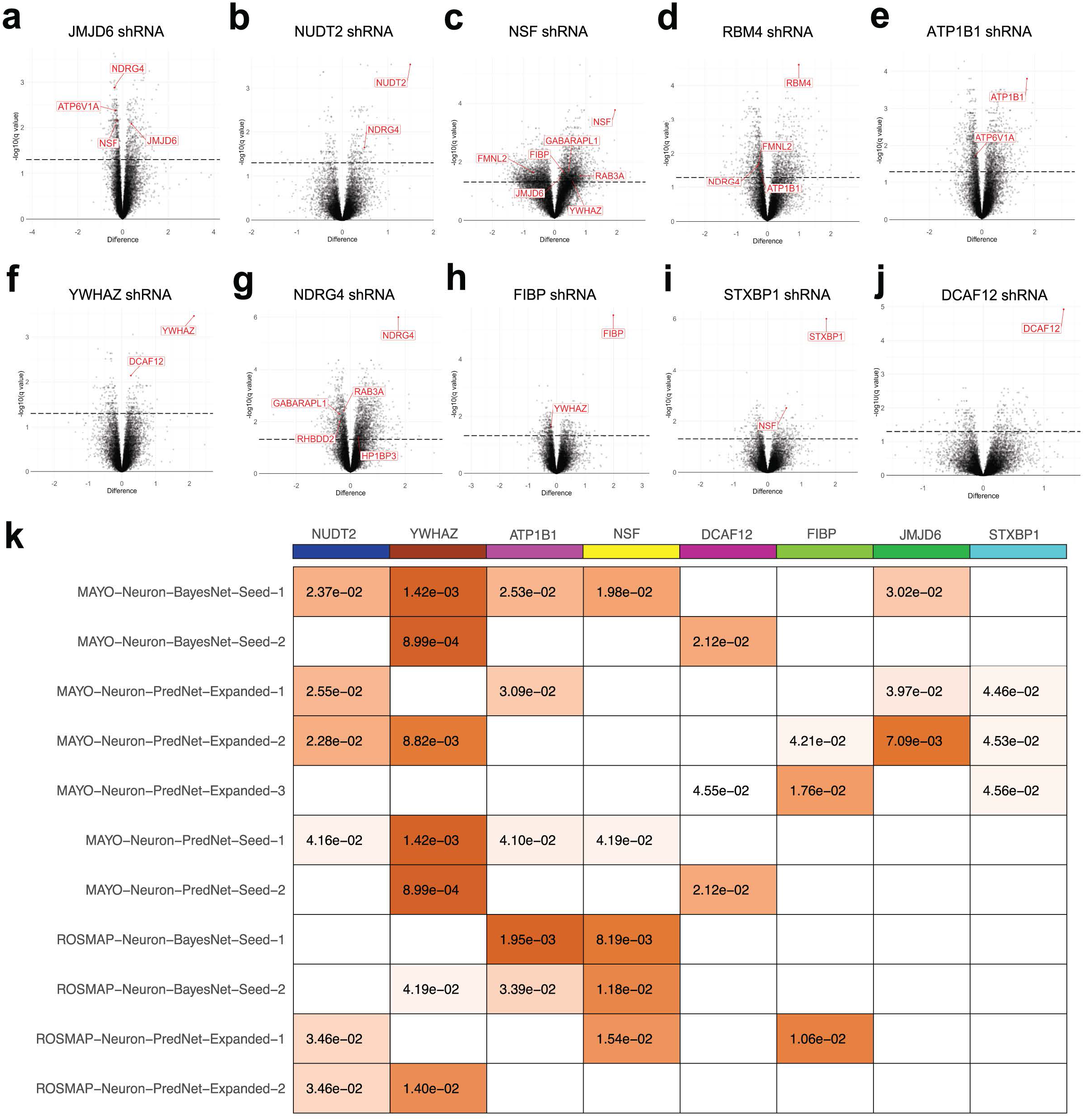
Gene expression changes following knockdown of the 10 validated targets in human iPSC-derived neurons. **(a-j)** RNAseq analysis showing significantly up- and down-regulated DE genes after shRNA-mediated knockdown of each of the 10 validated targets. Red gene symbols indicate any of our prioritized 19 targets genes that were significantly affected. Significance was assessed using the two-stage step-up method of Benjamini, Krieger, and Yekutieli with q-value<0.05, indicated by the black dashed line. (**k**) Network validation by enrichment analysis of significant DE genes following shRNA knockdown of the 10 validated targets in the 11 subnetwork networks. We compared the DE genes after knockdown of each target to the individual downstream subnetwork of that target extracted from the 11 reconstructed networks; significant enrichment is indicated in color with p-values (Fisher’s exact test, p<0.05). Detailed results are summarized in Supplementary Table S6.

We then further examined the gene expression changes resulting from knockdown of the 10 validated targets. Following *JMJD6* knockdown, which significantly altered ratios of both Aβ and tau in iNs, 656 genes were significantly upregulated and 419 genes significantly downregulated (Fig. 6a, q<0.05, determined using the two-stage linear step-up procedure of Benjamini, Krieger, and Yekutieli, with Q=5%). Interestingly, among those significantly upregulated genes were 3 of our other 19 key driver candidates (*NDRG4, ATP6V1A*, and *NSF*), indicating that their expression is affected by the reduction of *JMJD6* in neurons (Fig. 6a). Volcano plots in Fig. 6b-j highlight additional key driver candidates whose expression was affected by knockdown of each of the 9 other validated targets. Moreover, we found certain common genes affected by the perturbation of multiple validated targets: 6 genes (*FGF11, GIT2, KLHL28, PLCB3, SEPSECS*, and *SLC48A1*) were affected by knockdown of *NDRG4, STXBP1, YWHAZ*, and *JMJD6*, and 9 genes (*SEPTIN3, ABR, AOC2, CTFIP2, ZGTF2H1, MRPL17, NIIPSNAP1*, *RIMS4*, and *TMEM246*) were affected by perturbation of *DCAF12, NSF*, and *NUDT2*. This observation indicates that there may be unique and common molecular pathways among these validated AD endophenotype modulating targets.

To investigate possible mechanisms underlying these observations, we extracted regulatory pathways among the 10 validated targets in each of the 11 MAYO- and ROSMAP-neuron networks. We found that these 10 targets tightly regulate each other, and, interestingly, are all upstream regulators of the prominent proteins REST and VGF (Fig. 7a,b). REST (restrictive element 1-silencing transcription factor) is a known master regulator of neurogenesis via epigenetic mechanisms, apoptosis, and oxidative stress[126, 127]; VGF is a recently identified AD target whose overexpression in a mouse model reversed AD phenotypes[128]. In particular, our networks identified *FIBP* as a direct upstream regulator of VGF. Our findings thus indicate that these 10 targets may modulate AD-related pathology partially through REST and VGF pathways (Fig. 7b).

**Figure 7.**
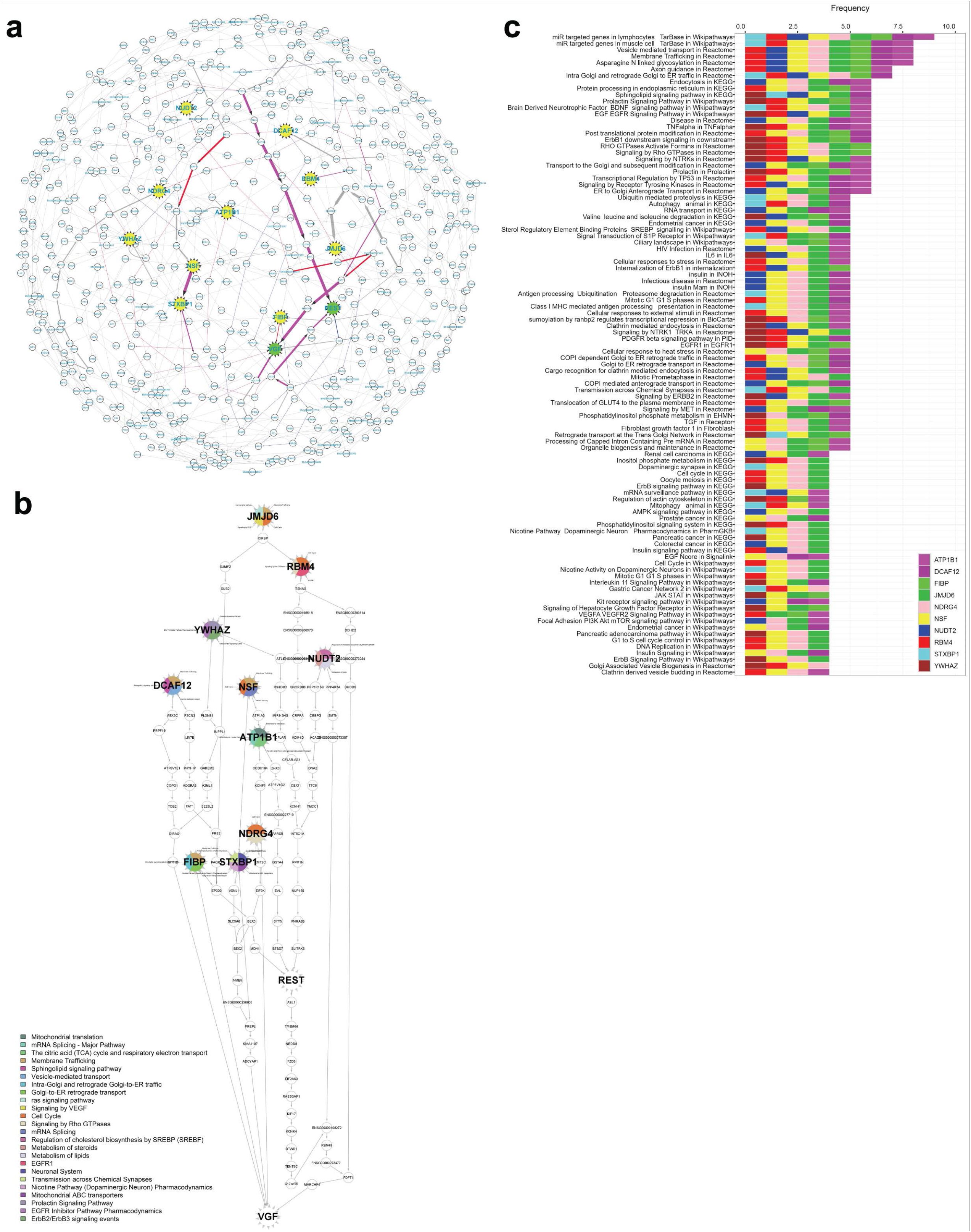
Regulatory pathway analysis reveals unique and shared biological pathways between the validated targets in each network. (**a**) The downstream members of each validated target were extracted from every network model, and edges from each downstream sub-network were pooled together into a consensus subnetwork of the 10 validated targets. As indicated by colored pathways, the 10 targets (yellow nodes) tightly regulate each other and are upstream regulators of both REST and VGF (green nodes). Edge thickness indicates the frequency of corresponding edges appearing across all networks. (**b**) The shortest paths from each of the 10 targets to REST and VGF were extracted from each network and pooled together into a hierarchical structure. The coloring of each target node annotates its representative enriched ConsensusPathDB (CPDB) pathways by significant DE genes in the shRNA knockdown experiments, as further detailed in Supplementary Table S7. (**c**) The overall CPDB pathways significantly enriched by each of the 10 target genes were pooled and ranked in descending order by the frequency of enrichment by any of the targets. Significance was assessed by Fisher’s exact test with p-value<0.05.

Finally, we performed pathway enrichment analysis (Methods) on the DE signatures derived from the RNAseq data in order to identify the unique and shared pathways affected by the knockdown of the 10 AD endophenotype modulating targets (*JMJD6, NSF, NUDT2, DCAF12, RBM4, YWHAZ, NDRG4, STXBP1, FIBP* and *ATP1B1*). We compared significant pathways enriched by the DE signature of each of the targets and found that 1 pathway is shared by 9/10 targets, 4 pathways are shared by 8/10 targets, 2 pathways are shared by 7/10 targets, 18 pathways are shared by 6/10 targets, and 40 pathways are shared by 5/10 targets (Fig. 7c, p-value<0.05). Comprehensive descriptions of all pathways affected by these targets are included in Supplementary Table S7.

## Discussion

Alzheimer’s disease (AD) is the most common neurodegenerative disease in the world, affecting millions of people worldwide. In the United States alone, an estimated 5.8 million Americans are currently living with AD dementia and this number is anticipated to reach 13.8 million by 2025[129]. Previous studies of LOAD pathogenesis using multi-omic data have identified numerous targets [19–22, 128]. However, although neurons are the principal cell type affected by AD etiology, the molecular mechanisms and therapeutic targets for AD revealed by these studies are not specific to neurons due to a lack of large-scale single-cell RNAseq data on neurons in AD. Thus, a comprehensive characterization of neuron-specific gene regulatory networks with association to AD is crucial to provide insight into the underlying causes of Alzheimer’s disease.

In this study, we employed a self-developed computational systems biology approach to model AD neuronal genetic regulation networks, with which we identified upstream regulators (key drivers) in neurons that contribute to AD pathology. In our pipeline, we employed PSEA to deconvolute RNAseq data from brain region-specific tissue in the MAYO and ROSMAP cohorts into five major cell types in the CNS including neurons, microglia, astrocytes, endothelial cells, and oligodendrocytes. In this study, we focused on the neuron-specific gene expression data and performed basic bioinformatics analyses including differential expression (DE) analysis, eQTL identification, co-expression module networks, and pathway enrichment analysis, followed by construction of causal network models and key driver gene identification.

From the network models, we identified a total of 1,563 neuronal key drivers which may represent new therapeutic targets. We used an unbiased ranking approach to prioritize 19 predicted key drivers for *in vitro* experimentation and tested the effects of their knockdown on the central components of the pathological hallmarks of AD, amyloid-β peptides (Aβ38, Aβ40, Aβ42) and phosphorylated tau protein, in a human iN system. We validated 10 targets which affected Aβ (*JMJD6, NSF, NUDT2, DCAF12, RBM4, YWHAZ, NDRG4*, and *STXBP1*) and/or tau/p-tau levels (*JMJD6, FIBP*, and *ATP1B1*). Only *YWHAZ* has been previously linked to AD through expression and mechanistic studies[130–134], while others have not yet been studied. Our findings of alterations to the neurotoxic ratios of both Aβ42 to Aβ40 and p231-tau to tau suggest therapeutic potential to both early and later stages of disease considering known patterns of pathology development in AD[135].

Most interestingly, we identified that knockdown of *JMJD6* (Jumonji Domain Containing 6, Arginine Demethylase and Lysine Hydroxylase) significantly increased both Aβ42 to 40 and p231-tau to tau ratios, suggesting therapeutic relevance to multiple stages of AD pathology. *JMJD6* belongs to the JmjC domain-containing family, catalyzes protein hydroxylation and histone demethylation, and appears to interact with distinct molecular pathways through epigenetic modifications of the genome[136, 137]. It is expressed in many tissues throughout the body, including the brain according to the Human Protein Atlas[138], but very little is known about its role in the brain or in neurodegenerative disease. However, based on its known role in epigenetic regulation epigenetic, it is expected that reduction of *JMJD6* expression may result in widespread changes in gene expression. Indeed, consistent with this prediction, we observed expression changes in a large number of genes following neuronal knockdown of *JMJD6*, including alteration of the expression of 3 other key driver targets of interest highlighted in this study (*NDRG4, ATP6V1A*, and *NSF*). Moreover, we found that an interesting association between *JMJD6* (as well as *NUDT2* and *NDRG4* among the 10 validated targets) and allele dosage. These 3 key driver genes are significantly associated with single nucleotide polymorphisms (SNPs) in their promoter regions (*cis*-eQTLs) in the MAYO and ROSMAP cohorts, further indicating that these genes may be actionable targets for AD therapeutic development.

We recognize that one caveat of our experimental system is that the neurons in a dish are not the same as neurons present in the aged AD brain; however, they do represent a powerful system for interrogating molecular connections between gene expression and proteins relevant to AD (namely, Aβ and tau). In our recent study, we showed that neurons derived from >50 different individuals show concordance between their levels of specific Aβ peptides and p-tau species and levels of these same proteins expressed in the brains of the same individuals[57]. Further, we showed concordance between protein and RNA module expressions between the iPSC-derived neurons and the brain tissue of the same people. Taken together, these results suggest that in spite of the reductionist nature of the system and the lack of aging, molecular networks are captured within the cells *in vitro* that are reflected in changes in Aβ and tau. Here, we employ this same experimental system to show that targeted reduction of *JMJD6* levels in human neurons induces effects on Aβ ratios and tau levels and phosphorylation.

Through our network models, we also discovered two shared downstream effectors of the 10 validated targets, which potentially explain the observed modulation of AD pathology: REST (restrictive element 1-silencing transcription factor) and VGF (VGF nerve growth factor inducible). REST is a known master regulator of neurogenesis via epigenetic mechanisms, apoptosis, and oxidative stress [126, 127] whose loss has been causally linked to Alzheimer’s disease [139, 140]. Additionally, recent studies have identified an association between changes in the epigenome, such as DNA methylation and histone modification, with changes in cognitive functions such as learning and memory[141–150]. Thus, dysregulation of epigenetic mechanisms through modulation of the targets may play a role in the pathogenesis of AD [142, 151]. VGF is also a target of interest which was recently validated to partially rescue memory impairment and neuropathology in 5xFAD mice[128]. Overexpression of VGF increased levels of activated BDNF receptor and adult hippocampal neurogenesis, which in turn regulated postsynaptic protein PSD-95 and improved cognition in the 5xFAD mice[128]. Pathway enrichment analysis confirmed that all 10 key drivers and their downstream genes in the network models were also significantly enriched for a variety of convergent and unique downstream cellular processes and functions which may explain additional molecular mechanisms at play, including vesicle-mediated membrane trafficking (common downstream of 8 targets); axon guidance, intra-Golgi trafficking, and retrograde Golgi-to-ER trafficking (common of 7 targets); and signaling pathways for sphingolipids, prolactin, BDNF/NTRKs, EGF-EGFR, TNFα, RHO GTPases, TP53, receptor tyrosine kinases (RTKs), and ER-to-Golgi transport (common of 6 targets).

In summary, our innovative computational systems biology approach using predictive network modeling has identified 10 targets which significantly modulate AD pathology via regulation of a variety of downstream pathways. These processes involve a wide spectrum of cellular pathways and possible mechanisms, and our results offer novel insights into potential therapeutic targets for drug discovery in Alzheimer’s disease.

## Acknowledgements

The authors thank the following funding resources which supported this study: NIH/NIA 1R56AG062620-01 and NIH/NINDS/Mayo Clinic U54NS110435 subaward to R.C.; NIH/NIA 1RF1AG057457-01 to R.C. and T.L.Y.-P.; and NIH/NIA R01AG055909 and RF1NS117446 to T.L.Y.-P.

R.C. is the founder of INTelico Therapeutics LLC and a co-founder of PATH Biotech LLC. This study is not supported by any funding from INTelico Therapeutics LLC or PATH Biotech LLC.

## Author Contributions

Conceptualization: T.L.Y.-P. and R.C.

ROSMAP RNAseq and WGS pre-processing: M.Y.R.H.

MAYO RNAseq and WGS pre-processing: M.Y.R.H., N.E.T.

Single cell-type gene expression: K.Z., R.C., M.Y.R.H., N.E.T.

Data analysis: K.Z., B.L., S.M., M.A., M.Y.R.H., R.C.

iNs shRNA experiment design and implementation: J.P.M., R.V.P.II, T.L.Y.-P.

Technical assistance: D.B.

Manuscript writing and figures: K.Z., J.P.M., R.C.

Manuscript editing: R.C., T.L.Y.-P.

## Methods

### 1. AMP-AD Consortium Data Source

Data was downloaded from the Accelerating Medicines Partnership – Alzheimer’s Disease (AMP-AD) consortium database hosted on the Synapse.org data portal (doi:10.7303/syn2580853).

### 2. AMP-AD Mayo Clinic Cohort and Data Pre-Processing

#### 2.1) Mayo Clinic Transcriptome and Genome-Wide Genotype Data

The Mayo Clinic (herein MAYO) transcriptome and genome-wide genotype datasets utilized in this study have previously been described[152–155]. The MAYO temporal cortex (TCX) RNA sequencing (RNAseq) data (Synapse ID: syn3163039) and genome-wide genotype data (Synapse ID: syn8650953) are available on the AMP-AD Knowledge Portal. We provide details on these datasets below.

#### 2.2) MAYO Cohort Participants

The overall MAYO dataset includes 278 subjects with the following diagnoses: 84 Alzheimer’s disease (AD), 84 progressive supranuclear palsy (PSP), 80 cognitively normal (CN) controls, and 30 pathologic aging. Subjects with AD each had a definite neuropathologic diagnosis according to the NINCDS-ADRDA criteria[156] and a Braak[157] neurofibrillary tangle (NFT) stage of ≥4.0. Control subjects each had a Braak NFT stage of 3.0 or less and CERAD[158] neuritic and cortical plaque densities of 0 (none) or 1 (sparse), and each lacked any of the following pathologic diagnoses: AD, Parkinson’s disease (PD), dementia with Lewy bodies (DLB), vascular dementia (VaD), progressive supranuclear palsy (PSP), motor neuron disease (MND), corticobasal degeneration (CBD), Pick’s disease (PiD), Huntington’s disease (HD), frontotemporal lobar degeneration (FTLD), hippocampal sclerosis (HipScl), or dementia lacking distinctive histology (DLDH). In the MAYO dataset, all disease subjects had ages at death ≥60 years; a more relaxed age cutoff of ≥50 years was applied for CN controls to achieve a sample size similar to that of the AD subjects, but we note there were only two additional control subjects with age at death below 60. This work was approved by the Mayo Clinic Institutional Review Board. All human subjects or their next of kin provided informed consent.

#### 2.3) MAYO RNAseq Data

TCX samples from all MAYO subjects underwent RNA extraction via the TRIzol/chloroform/ethanol method, followed by DNase treatment and cleanup of RNA using Qiagen’s RNase-Free DNase Set and RNeasy Mini Kit (Germantown, MD). Quantity and quality of all RNA samples were determined using the Agilent RNA 6000 Nano Kit on the Agilent 2100 Bioanalyzer system (Agilent Technologies, Santa Clara, CA). Only samples with an RNA Integrity Number (RIN) ≥5.0 were included in this study. MAYO RNAseq samples were randomized across flowcells, taking into account age at death, sex, RIN, Braak stage, and diagnosis. Library preparation and sequencing of the samples were conducted at the Mayo Clinic Medical Genome Facility Genome Analysis Core, as previously described[159]. The TruSeq RNA Sample Prep Kit (Illumina) was used for library preparation from all samples. Library concentration and size distribution were determined using an Agilent Bioanalyzer DNA 1000 Kit (Agilent Technologies). Three samples were run per flowcell lane using barcoding. All samples underwent 101 base-pair (bp), paired-end sequencing on Illumina HiSeq2000 instruments. Base-calling was performed using Illumina’s Real-Time Analysis (RTA) 1.17.21.3. FASTQ sequence reads were aligned to the human reference genome using TopHat 2.0.12[160] and Bowtie 1.1.0[161], and Subread 1.4.4 was used for gene counting[162]. FastQC[163] was used for quality control (QC) of raw sequence reads, and RSeQC[164] was used for QC of mapped reads.

#### 2.4) MAYO RNAseq Data Processing and Quality Control

All MAYO RNAseq samples had percentage of mapped reads ≥85%. In the first step, we used R statistical software (R Foundation for Statistical Computing, version 3.2.3) to transform the raw read counts to counts per million (CPM), which were then log2-normalized. Mean expression levels of Y chromosome genes with non-zero counts were plotted to identify any samples with deviation from expected expression based on recorded sex; 2 AD samples were identified with discordant sex and excluded. In the second step, raw read counts were normalized using Conditional Quantile Normalization (CQN) via the Bioconductor package[165], accounting for sequencing depth, gene length, and GC content. Before running CQN normalization, GC content was calculated via Repitools in the Bioconductor package[166] and sequencing depth was calculated as the sum of reads mapped to genes. Genes with non-zero counts across all samples were retained and principal component analysis (PCA) was performed using the prcomp function implemented with R statistical software. Principal components 1 and 2 were plotted and no outliers (>6 SD from mean) were identified. We additionally excluded 2 samples due to missing data in one or more key covariates.

#### 2.5) MAYO Genome-Wide Genotype Data

Subjects in the MAYO RNAseq cohort underwent whole genome genotyping using the Illumina Infinium HumanOmni2.5-8 BeadChip Kit (San Diego, CA), which delivers comprehensive coverage of both common and rare single nucleotide polymorphism (SNP) content from the 1000 Genomes Project[167] (minor allele frequency >2.5%) and provides genotypes for 2,338,671 markers. Genotyping was performed at the Mayo Clinic Medical Genome Facility Genome Analysis Core. Whole genome genotype calls were made using the auto-calling algorithm in Illumina’s BeadStudio 2.0 software, after which they were converted into PLINK formats for analysis[168].

#### 2.6) MAYO Genotype Data Quality Control

All genome-wide genotype samples were checked for discordant sex; the same 2 AD subjects excluded for this reason in Section 2.4 were identified. Subjects were assessed for heterozygosity rates >3 SD from the mean. One AD sample had high heterozygosity with respect to the mean, indicating possible sample contamination, and 3 samples (2 controls and 1 AD) had low heterozygosity with respect to the mean, indicating either divergent ancestry or consanguinity; these 4 samples were also excluded from the analysis. The dataset was then filtered to include only autosomal SNPs. PLINK was used to identify any sample duplicates or related pairs of subjects. Two pairs of samples were identified as >3^rd^ degree relatives; for each pair, the sample with the lower SNP call rate was excluded. The dataset was further filtered to remove complex genomic regions (chr8:1-12,700,000; chr2:129,900,001-136,800,000; chr17:40,900,001-44,900,000; and chr6:32,100,001-33,500,000) and linkage disequilibrium (LD) pruned using the SNPRelate (v1.4.2) package in R (v3.2.3) [169], implementing an LD threshold of 0.15 and a sliding window of 1E-07 bp. Remaining SNPs and subjects were analyzed using EIGENSOFT[170] for population outliers. Two samples were identified as population outliers using the default parameter of >6 SD from the mean on any of the top 10 inferred axes following 5 iterations, and they were removed from further analysis.

After QC of the MAYO RNAseq and genome-wide genotype datasets, we obtained a total of 266 subjects with matched transcriptome and genotype data, including 76 CN subjects and 79 AD subjects which were used in all subsequent analysis. We performed rigorous statistical testing to demonstrate that these samples are well balanced with respect to age at death (p-value=0.57) as well as sex (p-value=0.24) (Fig. S7a,c).

### 3. AMP-AD ROSMAP Cohort and Data Pre-Processing

#### 3.1) ROSMAP Transcriptome and Genome-Wide Genotype Data

The Religious Orders Study and Memory and Aging Project (herein ROSMAP) dataset dorsolateral prefrontal cortex (DLPFC) gene expression (RNAseq BAM files), genotypes, and clinical covariates were downloaded from the Synapse.org data portal (Synapse IDs for the respective data types: syn4164376, syn3157325, and syn3191087) using the synapseClient R library[171]. Requests for ROSMAP data can be made at https://www.radc.rush.edu/.

#### 3.2) ROSMAP Cohort Participants

The ROSMAP dataset contains two cohorts: the Religious Orders Study (ROS) and the Memory and Aging Project (MAP)[172]. Both ROS and MAP are longitudinal clinical-pathologic cohort studies of aging and dementia run by the Rush Alzheimer’s Disease Center in Chicago, IL. In both studies, all participants enroll without known dementia and agree to annual clinical evaluation and brain donation as a condition of entry. ROS has enrolled individuals from religious orders from across the United States starting in 1994, and MAP has enrolled lay persons from across northeastern Illinois since 1997. Each study annually administers a battery of 21 cognitive performance tests, 19 of which are in common. Alzheimer’s disease status is determined by a computer algorithm based on cognitive test performance with a series of discrete clinical judgments made by both a neuropsychologist and a clinician. First, subjects are categorized as not cognitively impaired (NCI, if diagnosed without dementia), mild cognitive impairment (MCI), or Alzheimer’s disease (AD). Diagnoses of dementia and AD conform to standard definitions[172]. Next, a clinician reviews all cases determined by the algorithm to render a diagnosis blinded to data collected in prior years. In addition to dementia, 5 other diagnoses are determined by this approach, including stroke, cognitive impairment due to stroke, parkinsonism, Parkinson’s disease, and depression. Most of these other diagnoses are determined by self-report. Upon death, a summary diagnosis is made by a neuropsychologist blinded to post-mortem assessment. The post-mortem neuropathologic evaluation performed includes a uniform structured assessment of AD pathology, cerebral infarcts, Lewy body disease, and the other pathologies common in aging and dementia (e.g., vascular dementia or frontotemporal dementia). The evaluation procedures follow those outlined by the pathologic dataset recommended by the National Alzheimer’s Disease Coordinating Center. Pathologic diagnoses of AD use NIA-Reagan and modified CERAD criteria[173], and the evaluation of neurofibrillary pathology uses Braak staging[174]. The ROS and MAP studies are both conducted by the same clinical and pathologic data collection teams, with extensive item-level harmonization allowing the data to be efficiently merged.

#### 3.3) ROSMAP Genotype Data and Quality Control

PLINK 2.0[175] was used to perform operations on the genotype files, and positions were converted from hg18 to hg19 (http://genome.ucsc.edu/cgi-bin/hgLiftOver). Picard[176] was used to sort the resulting genotype files, and samples were removed using PLINK 2.0 if they had variants with >2% missing values, minor allele frequency <1%, Hardy-Weinberg equilibrium<10E-6, or inbreeding coefficient >0.15. We started with 750,173 variants in 1,708 individuals; after quality control, 736,073 variants in 1,091 individuals remained.

#### 3.4) ROSMAP RNAseq Data Processing

Out of the 1,091 ROSMAP subjects remaining after genotype QC, we obtained a total of 612 subjects which had matched RNAseq data. The RNA-seq BAM files were sorted using samtools[177] and converted to FASTQ files using the SamToFastq function[176]. RAPiD[178] was used to generate a count matrix for the gene expression data and a vcf file for each sample aligned to hg19 from the FASTQ files. ROSMAP RNAseq read count expression data was normalized using log2 counts per million (CPM) and the TMM method[179] implemented in edgeR[180]. Genes with over 1 CPM in at least 30% of the experiments were retained. We then used precision weights as implemented in the voom function from the limma[181] R package to further normalize the gene counts.

The total 612 subjects with matched QC data included 194 CN subjects and 212 AD subjects which were used in all subsequent analysis. Regarding the ROSMAP cohort, it has been noted that the range of age at death is broad but restricted to the older segment of the age distribution of the North American population and that age and sex are important confounders when performing any analyses of ROS and MAP data[76]. We observed this variance in the age at death (p-value<0.05) but found no significant difference in sex among the ROSMAP subjects used in our analysis (p-value=0.072) (Fig. S7b,d). To address the imbalanced age distribution, we later performed covariate adjustment for age (together with other covariates, Section 5), and we confirmed removal of the effects of age and other confounding variables by variance partition analysis (VPA) before and after covariate adjustment (Fig. S2 b,d).

### 4. MAYO and ROSMAP Genotype Data Imputation

After quality control of both datasets, we used 1000 Genomes Project[167] data and IMPUTEv2[182] to impute untyped variants. Imputed variants were removed if they failed any of the previously listed quality control criteria or had information scores <0.6. After imputation, we had 7,132,687 variants in MAYO and 9,333,139 variants in ROSMAP.

### 5. Deconvolution of RNAseq Data into Neuron-Specific Expression Residuals

After normalizing both MAYO and ROSMAP expression data (described above), expression residuals were obtained for each dataset respectively by adjusting for covariates using the limma R package[181]. For MAYO, expression residuals were obtained by correcting for the effects of technical confounding factors (i.e., sequencing batch), sample-specific variables (RNA integrity number [RIN], exonic mapping rate, source of tissue), and patient-specific covariates (sex, age at death, *APOE* genotype). For ROSMAP, we adjusted for a slightly different set of covariates due to a greater number of recorded measurements available: study (ROS or MAP), sequencing batch, post-mortem interval (PMI), RIN, exonic mapping rate, sex, educational attainment, *APOE* genotype, and age at death. For both MAYO and ROSMAP data, we computed the exonic mapping rate using RNAseQC[183].

In addition to the covariate adjustments listed above for both datasets, we also adjusted for five cell type markers[72]: *ENO2* [neuronal], *CD68* [microglial], *CD34* [endothelial], *OLIG2* [oligodendrocytic], and *GFAP* [astrocytic]. To obtain expression residuals that mimic expression patterns seen in neurons, for every gene, we added the *ENO2* effects estimated by the linear regression models back to the expression residuals. Comparing the variance of normalized gene expression before and after covariate adjustment, we confirmed removal of the effects from confounding variables (Fig. S2), allowing us to conclude that the residual results are unbiased and robust against these adjusted covariates.

The final neuron-specific expression residual data available for further analysis included 19,885 genes from 155 individuals in MAYO (76 CN, 79 AD) and 20,276 genes from and 406 individuals in ROSMAP (194 CN, 212 AD), with 18,408 genes common to both datasets which are comparable with processed residuals of the same cohorts on AMP-AD knowledge portal[184].

#### 5.1) Rationalization and Validation of Single-Gene Biomarkers for Bulk-Tissue RNAseq Deconvolution

Our rationale for using single-gene biomarkers over multi-gene biomarkers derived from singlecell RNAseq (scRNAseq) data was manifold. First, multi-gene biomarkers derived from various scRNAseq studies in control human brains [47–51] show no significant overlap among themselves, indicating a lack of robustness and consensus in these biomarkers derived from scRNAseq studies (Fig. S3a). Second, PCA analysis shows a prominent overlap of scRNAseq biomarker expression across different CNS cell types in MAYO and ROSMAP AD data, indicating that the majority of scRNAseq-derived biomarker gene expression is convoluted and reflecting potential interactions between different cell types under the AD condition (Fig. S3b,c). Furthermore, there is significant overlap between scRNAseq-derived biomarkers and AD therapeutic targets in the AMP-AD Agora knowledge portal[121] (Fig. S3d,e); this overlap is more significant than randomly selected genes from the background overlapping with the Agora list, indicating that scRNAseq-derived biomarkers may play a significant role in AD pathology. For these reasons, multi-gene biomarkers of cell type are not ideal for adjusting the bulk-tissue gene expression variance by population-specific expression analysis (PSEA). By contrast, our singlegene biomarkers are derived from biological knowledge and have been validated by other groups [72]. Moreover, our single-gene biomarkers had no overlap with AD therapeutic targets in the Agora list, thus making them good candidates for PSEA. Lastly, our neuron-specific residual derived from single-gene biomarkers is significantly correlated with the “pseudo” neuron-specific residuals derived from a randomly selected subset of scRNAseq biomarkers by PSEA (Fig. S4), indicating that our neuron-specific residual represents a robust neuronal component in the bulk-tissue RNAseq data for neuron-specific therapeutic target discovery in LOAD. Specific methods behind the analyses described above are elaborated in the following subsections.

#### 5.2) Comparison of scRNAseq-Derived Biomarker Genes Per Cell Type

We assembled cell-type specific biomarkers derived from existing scRNAseq studies for neurons, microglia, astrocytes, endothelial cells, and oligodendrocytes. These biomarkers are listed in Supplementary Table S8. For each cell type, we compared the biomarker genes by pairing each scRNAseq study and calculating the significance of the overlap by Fisher’s exact test (Fig. S3a; FDR>0.05, 1/0/1/1 significant pairs out of 6 study-pairs in astrocytic/endothelial/microglial/oligodendrocytic types and 2 significant pairs out of 10 study-pairs in neuronal types).

#### 5.3) PCA Analysis of scRNAseq-Derived Biomarker Expression in AMP-AD Data

After creating a merged biomarker list from different scRNAseq studies for each cell type as described, we then extracted the gene expression matrix of the merged biomarkers from the MAYO and ROSMAP RNAseq data and applied principal component analysis (PCA) on the extracted RNAseq sub-matrix (Fig. S3b,c).

#### 5.4) Comparison with AMP-AD Agora Targets

We also merged all biomarkers for each cell type and calculated the percentage of overlap with targets in the AMP-AD Agora list (Fig. S3d,e, see above). To evaluate the significance of this overlap, for each cell type, we simulated a background distribution of overlap by randomly selecting the same number (as on the merged biomarker list) of background genes – taking the non-duplicate union of genes in MAYO and ROSMAP RNAseq data – and comparing the randomly generated “pseudo” biomarker list to the AMP-AD Agora targets to generate an overlapping percentage. We repeated the random simulation 10,000 times to construct the background distribution. The p-value was then calculated per cell type by comparing the true percentage to the background distribution.

#### 5.5) Evaluation of Robustness of Deconvoluted Neuron-Specific Residuals

By using population-specific expression analysis (PSEA)[44], we estimated the variance component of the MAYO and ROSMAP bulk-tissue RNAseq data explained by our single-gene neuronal biomarker (*ENO2*). Next, we randomly selected a subset of biomarkers of each cell type from the scRNAseq-derived biomarkers (Supplementary Table S8), then again applied PSEA to estimate the variance component of the bulk-tissue RNAseq data explained by the simulated biomarker subset. We then calculated the Pearson correlation of each gene in the residual between our single-gene neuronal residuals and the simulated neuronal residuals. We repeated this procedure 1,000 times to construct a distribution of the correlations (Fig. S4).

Next, we constructed a background distribution of correlation. To this end, we again randomly selected a subset of “pseudo” biomarkers of each cell type from the background genes (the nonduplicate union of genes in MAYO and ROSMAP RNAseq data), then applied PSEA to estimate the variance component of the bulk-tissue RNAseq data explained by the simulated “pseudo” biomarkers. We then calculated the Pearson correlation of each gene between our single-gene neuronal residuals and the simulated “pseudo” neuronal residuals. We repeated this procedure 1,000 times to construct a distribution of the correlations (Fig. S4). Lastly, we applied a t-test to compare the correlation distribution described above with the background distribution described here (p-value<2.2E-16).

### 6. Computational Analysis of Neuron-Specific Gene Expression Data

#### 6.1) eQTL Analysis

Expression quantitative trait locus (eQTL) analysis was performed using the R package MatrixEQTL v2.1.1[185] using QCed genotypes and normalized and covariate-adjusted cell-type specific expression residuals. *cis*-eQTL analysis considered markers within 1 Mb of the transcription start site of each gene. False discovery rates (FDR) were computed using the Benjamini–Hochberg procedure[186].

#### 6.2) Differential Expression (DE) Analysis

Using linear models, as implemented in the limma R package [181], we interrogated the cell-type specific residual expression data for genes differentially expressed between AD cases and healthy controls. Significance was assessed using FDR<0.05.

#### 6.3) Pathway Enrichment Analysis

We downloaded pathways from ConsensusPathDataBase (CPDB)[83]. Given a set of genes, we performed enrichment analysis of each pathway over this set of genes by Fisher’s exact test.

#### 6.4) Co-Expression Network Analysis

Co-expression networks were constructed using the coexpp R package[187]. A soft thresholding parameter value of 6.5 was used to power the expression correlations. Seeding gene lists for the predictive networks were obtained by selecting genes in co-expression modules that were statistically enriched (FDR<0.05) for DE genes or neuronal cell markers[50].

#### 6.5) Key Driver Analysis

To perform Key Driver Analysis (KDA), we used the KDA R package [70] (KDA R package version 0.1, available at http://research.mssm.edu/multiscalenetwork/Resources.html). The package first defines a background sub-network by looking for a neighborhood k-step away from each node in the target gene list in the network. Then, stemming from each node in this sub-network, it assesses the enrichment in its k-step (k varies from 1 to K) downstream neighborhood for the target gene list. In this analysis, we used K = 6.

#### 6.6) Key Driver Prioritization

##### A) Impact Score

To rank the key drivers based on impact score, we calculated the following metrics:

1. KDA_unit_score. For each network, we performed KDA. We defined different target gene lists for KDA with several gene sets: i) the overlap between DE genes and selected modules, recording the number of overlapping gene sets nominating a given gene as a key driver; ii) the selected modules, recording the number of modules nominating a given gene as key driver; and iii) the DE gene set, whose value was 1 or 0, indicating whether (1) or not (0) DE genes nominated a given gene as a key driver.
2. KDA_score_sum. We summed the 3 KDA_unit_score values described in 1).
3. KDA_position_sum. We found the non-zero sum of the frequency of the 3 values in 1).
4. Normalized_priority_score. For each network, we sorted the key drivers first according to KDA_position_sum in descending order and then according to KDA_score_sum in descending order. In each network, we then calculated the normalized_priority_score by dividing each key driver’s rank by the maximum rank for the corresponding network.
5. Replication_count. We recorded the number of networks from which a key driver was derived.
6. Avg_priority_score. We calculated the averaged priority score per key driver across all networks by dividing normalized_priority_score by replication_count.
7. Avg_DE_R. Since a gene could appear in multiple KDA files and network files (e.g. gene A appearing in 5 KDA files and 7 networks), we selected each corresponding network from which a given key driver was derived, retrieved all downstream members in the networks of the key driver, and calculated the following metrics: i) DE_reach, the percentage of corresponding DE genes covered by the gene’s downstream effectors, describing an overall overlap between downstream effectors of a key driver and DE genes; ii) the number of DE genes (N) and a modified version of count (R) for each layer of a downstream subnetwork, where the modified count (R) was computed as following: for a given DE gene, if it had X parent nodes and among those X parents, Y of them were not downstream members of the given key driver, we would say the burden factor of the DE gene for the key driver is Y/X. Thus, for each layer in the downstream sub-network, if the layer contained Z DE genes, then N=Z and R=Z minus the sum of burden factors over all Z DE genes for the layer; iii) R.coeff, the coefficient of a linear model, where the response variable is a vector of the normalized (by number of layers) cumulative sum of R for each layer, thus describing the cumulative percentage of parents that are also DE genes between the first layer and the current layer (value between 0 and 1), and the predictor variable is the layer index. The higher this coefficient, the more impact a key driver has on the downstream DE genes; iv) impact_per_network score of a key driver, which was calculated as DE_reach*R.coeff and is a joint descriptor of the overall downstream-DE gene overlap and local DE percentage of every layer in the downstream sub-network of each key driver. Note that the above DE_reach and R.coeff values are calculated per network; and finally v) Avg_DE_R, the averaged value of impact_per_network for a key driver over all the networks to find its overall impact across all networks, since individual key drivers may be identified by multiple networks.
8. Impact_score. Finally, the impact score of a key driver was calculated as (1-avg_priority_score)*Replication_count*Avg_DE_R. We ranked the key drivers in descending order.

##### B) Robustness Score

To rank the key drivers based on robustness score, we calculated the following metrics:

1. Dataset_Count. We calculated how many datasets, i.e., MAYO and ROSMAP cohorts, by which a key driver was replicated.
2. Geneset_Count. We calculated how many types of gene sets, i.e., expanded or seeding gene sets, by which a key driver was replicated.
3. Avg_DE_R score. This score was calculated in the same manner as described above (step 7 of Impact Score).
4. We ranked the key drivers according to robustness first by Dataset_Count in descending order, then ranked by Geneset_Count in descending order, and lastly ranked by Avg_DE_R score in descending order.

#### 6.7) Bayesian Networks, Predictive Networks, and Network Validation

Although the co-expression network modules capture highly co-regulated genes operating in coherent biological pathways, these modules do not reflect the probabilistic causal information needed to identify key driver genes.

Bayesian networks (BNs)[188] are a long-standing form of statistical network modeling used to reverse-engineer probabilistic causality among variables; with the development of high-throughput sequencing technology, BNs have been widely used to infer causal gene regulatory networks in different diseases [189–194]. Recent studies have applied BNs to infer molecular mechanisms and key drivers in Alzheimer’s disease[22, 128].

However, BNs have significant limitations with respect to inferring opposite causality given the symmetry of joint probability. Recent work has demonstrated that bottom-up causality inference can accurately distinguish true causality from opposite causality in equivalent classes[69]. In this study, we developed a novel computational network model, called predictive network modeling, by integrating conventional (top-down) Bayesian networks with bottom-up causality inference in order to address the problem of opposite causality inference in BN modeling. Here, our causal predictive network pipeline incorporates multi-scale omics data, including genotypes and transcriptomic profiles, in the MAYO and ROSMAP datasets (deconvoluted neuron-specific residuals) in order to build causal predictive networks separately in both datasets.

#### 6.8) PathFinder Pathway Analysis

The PathFinder method[58] is based on the classical Depth First Search (DFS) algorithm[195]. The goal of PathFinder is to expand the initial target gene set by including genes in the background network located in the paths connecting input genes.

Since the background network could contain directed and undirected edges, we transformed the undirected edges into two edges with the same two end nodes but different directions. We did not allow these two edges to form a loop and simultaneously appear in one path.

The DFS algorithm starts from one input gene and stops if the length of path it explores reaches K or if the path arrives at a node without a valid child node. Whenever any of the stop criteria above was satisfied, we checked whether the path contained at least two input genes. If not, the path was discarded. Otherwise, among all the input genes in the path, we determined the target gene with the maximum distance to the starting input gene, and all the nodes between this gene and the starting input gene were then included in the seeding gene list for the network. In practice, we ran DFS for each input gene and combined the results to get the final network seeding gene list.

### 7. Induced Pluripotent Stem Cell (iPSC) Derived Neuron Culture and Assays

#### 7.1) iPSC Maintenance and Induced Neuron Differentiation

The human control iPSC line YZ1 was obtained from the University of Connecticut Stem Cell Core facility and was maintained in StemFlex Medium (Thermo Fisher Scientific, Waltham, MA). Induced neurons (iNs) were generated as described [24, 30, 60], with minor modifications described below. Briefly, iPSCs were plated in mTeSR1 media (STEMCELL Technologies, Vancouver, Canada) at a density of 95K cells/cm^2^ on Matrigel-coated plates (Corning Inc., Corning, NY) for viral transduction. Media was changed from StemFlex to mTeSR1 as we found better transduction viability with mTeSR1. Viral plasmids were obtained from Addgene (plasmids #19780, 52047, 30130; Watertown, MA). FUdeltaGW-rtTA was a gift from Konrad Hochedlinger (Addgene plasmid #19780), and Tet-O-FUW-EGFP (Addgene plasmid #30130) and pTet-O-Ngn2-puro (Addgene plasmid #52047) were gifts from Marius Wernig. Lentiviruses were obtained from ALSTEM (Richmond, CA) with ultra-high titers and used at the following concentrations: pTet-O-NGN2-puro: 0.1 μl/50K cells; Tet-O-FUW-eGFP: 0.05μl/50K cells; Fudelta GW-rtTA: 0.11μl/50K cells. Transduced cells were dissociated with Accutase (Gibco, Thermo Fisher Scientific) and plated onto Matrigel-coated plates at 50,000 cells/cm^2^ in mTeSR1 (day 0). On day 1, media was changed to KSR media with doxycycline (2 μg/ml, Sigma-Aldrich, St. Louis, MO). Doxycycline was maintained in the media for the remainder of the differentiation. On day 2, media was changed to 1:1 KSR:N2B media with puromycin (5 μg/ml, Gibco). On day 3, media was changed to N2B media + 1:100 B27 supplement and puromycin (10 μg/ml). Puromycin was maintained at this concentration in the media for the remainder of the differentiation. From day 4 onwards, cells were cultured in NBM media + 1:50 B27 + BDNF, GDNF, CNTF (10 ng/ml each, PeproTech, Rocky Hill, NJ) + doxycycline and puromycin as described.

#### 7.2) Induced Neuron Media

- *KSR media*: Knockout DMEM (Gibco), 15% KOSR (Invitrogen, Thermo Fisher Scientific), 1x MEM-NEAA (Invitrogen), 55 μM beta-mercaptoethanol (Invitrogen), 1x GlutaMAX (Life Technologies, Thermo Fisher Scientific).
- *N2B media*: DMEM/F12 (Life Technologies), 1x GlutaMAX (Life Technologies), 1x N2 supplement B (STEMCELL Technologies), 0.3% dextrose (D-(+)-glucose, Sigma-Aldrich).
- *NBM media*: Neurobasal Medium (Gibco), 0.5x MEM-NEAA (Invitrogen), 1x GlutaMAX (Life Technologies), 0.3% dextrose (D-(+)-glucose, Sigma-Aldrich).

#### 7.3) Induced Neuron Lentiviral Transduction

At d17 of differentiation, neurons were transduced with lentiviruses encoding shRNA constructs against selected targets (Broad Institute, Cambridge, MA), as described in[20]. For each round of experiments, two controls were included: a lentivirus expressing the pLKO vector without an shRNA (“empty”) or else not transduced (fresh media only). iNs were transduced with a 1:1 ratio of media to lentivirus. Following ~18 hours of incubation, media containing virus was removed and replaced with fresh media, and cells were incubated for an additional 96 hours. On d22 of differentiation, conditioned media was then collected and stored at −20°C for Aβ analyses, and cells were lysed either for RNA purification or protein harvest. Gene knockdowns were confirmed by qPCR.

#### 7.4) Aβ ELISA

Aβ present in the conditioned media was measured by the 6E10 Aβ Peptide Panel Multiplex ELISA (Meso Scale Discovery, Rockville, MD) following manufacturer instructions. Briefly, conditioned media from transduced cells were incubated in pre-blocked wells along with detection antibody solution. Plates were read using an MSD SECTOR Imager 2400 and resulting peptide concentrations were normalized to total protein in the cell lysate per well measured using the Pierce BCA Protein Assay Kit (Thermo Fisher Scientific). Data for each shRNA knockdown were additionally normalized to the average of control conditions for each parameter measured.

#### 7.5) Tau ELISA

Protein was extracted from iNs by lysis in NP-40 lysis buffer (1% NP40, 0.5M EDTA, 5M NaCl, 1M Tris) containing cOmplete protease inhibitors and phosSTOP (Roche, Penzberg, Germany). Lysates were analyzed using the Multi-Spot Phospho (Thr 231)/Total Tau ELISA (Meso Scale Discovery) following manufacturer instructions. Briefly, lysates were incubated in pre-blocked wells for 1 hr prior to detection antibody application for 1 hr. Plates were read using an MSD SECTOR Imager 2400 and resulting concentrations were again normalized to total protein in the cell lysate per well (Pierce BCA Protein Assay Kit) and data for each shRNA knockdown were. normalized to the average of control concentrations for each parameter.

#### 7.6) Induced Neuron RNA sequencing

For iNs, at least 250 ng of total RNA input was oligo(dT) purified using the PureLink RNA Mini Kit (Invitrogen), then double-stranded cDNA was synthesized using SuperScript III Reverse Transcriptase (Invitrogen) with random hexamers. RIN >9 was confirmed using the Agilent 4200 TapeStation system (Agilent Technologies). RNAseq on the shRNA-treated iNs was performed by Functional Genomics Core at the University of Arizona at a depth of 30 million single-end reads (100 bp long). The RNAseq data was QCed and processed with the same steps as outlined in ROSMAP RNAseq Data Processing (section 3.4).

### 8. Statistical Analyses

All statistical analyses were performed in R unless otherwise noted.

## Supplementary Information

**Figure S1.**
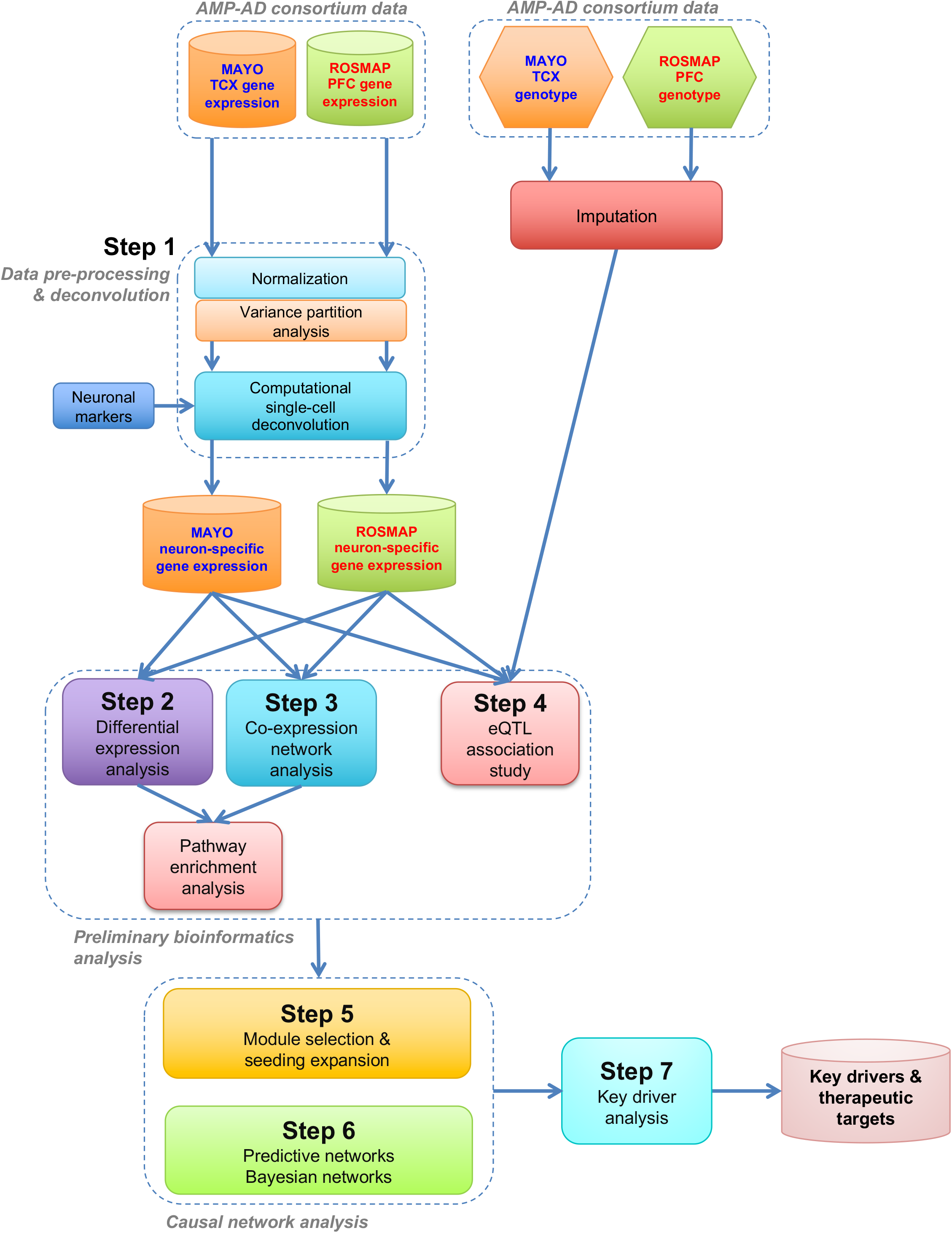
Workflow of our network analysis pipeline integrating two independent brain transcriptome and genome-wide genotype datasets (MAYO and ROSMAP) to construct neuron-specific predictive networks of AD and predict key drivers (therapeutic targets) associated with AD pathology. TCX: temporal cortex; PFC: prefrontal cortex.

**Figure S2.**
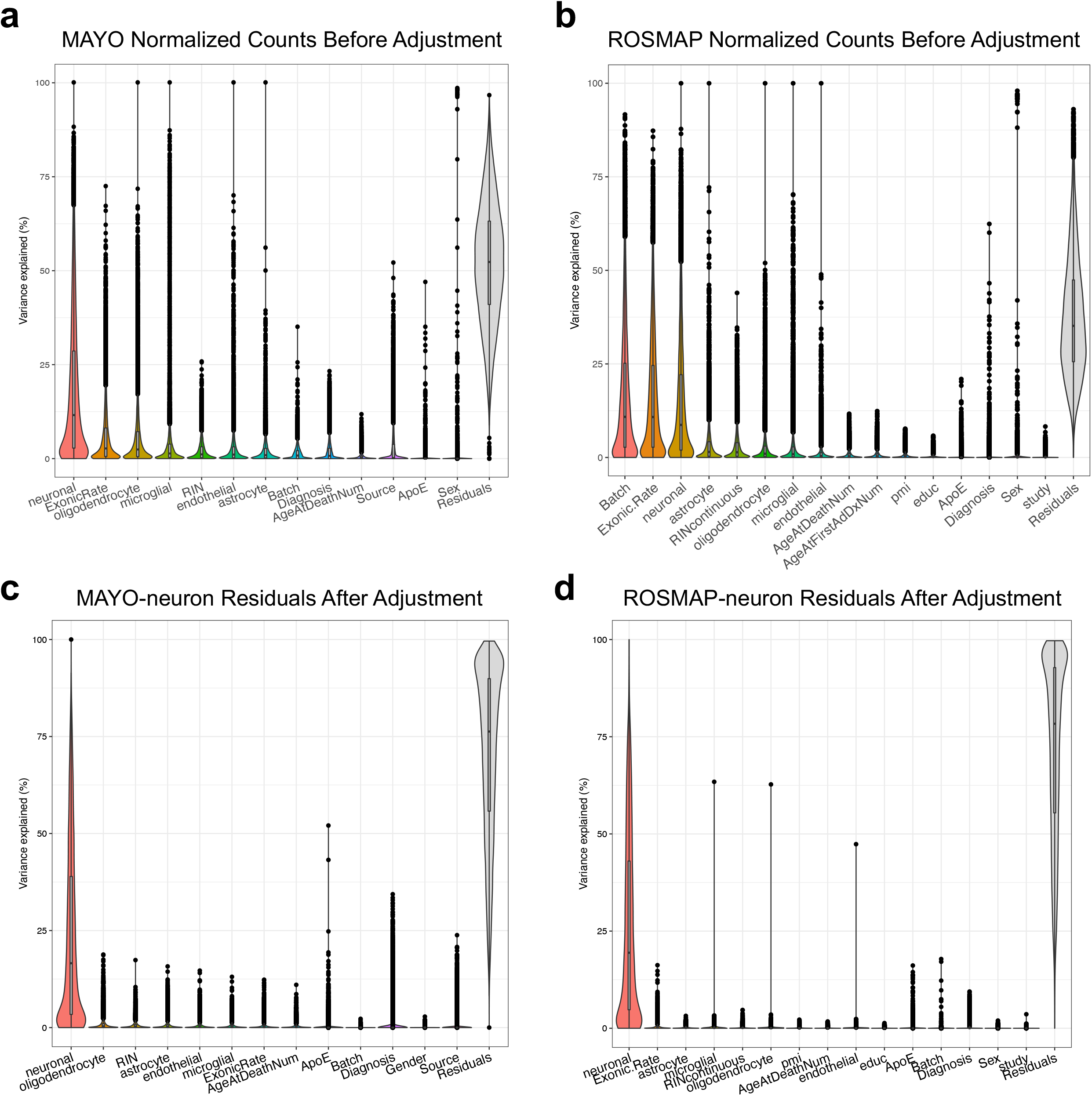
(**a,b**) Gene expression variance partition analysis (VPA) of bulk-tissue RNAseq data in the MAYO (**a**) and ROSMAP (**b**) cohorts *<u>before</u>* deconvolution and covariate adjustment reveals a prominent effect of cell type on bulk-tissue gene expression in the brain. *ENO2, CD68, GFAP, CD34*, and *OLIG2* were used as cell-type specific marker genes for neurons, microglia, astrocytes, endothelial cells, and oligodendrocytes, respectively. ExonicRate: exonic mapping rate; RIN or RINcontinuous: RNA integrity number; AgeAtFirstADDxNum: age at first AD diagnosis; pmi: post-mortem interval; educ: education. (**c,d**) The variance partition analysis on neuron-specific gene expression residuals in MAYO (**c**) and ROSMAP (**d**) *<u>after</u>* de-convolution and covariate adjustment demonstrates that the neuron-specific residuals capture the neuronal component (variance) and that the effects of other covariates and other cell types in the brain are removed.

**Figure S3.**
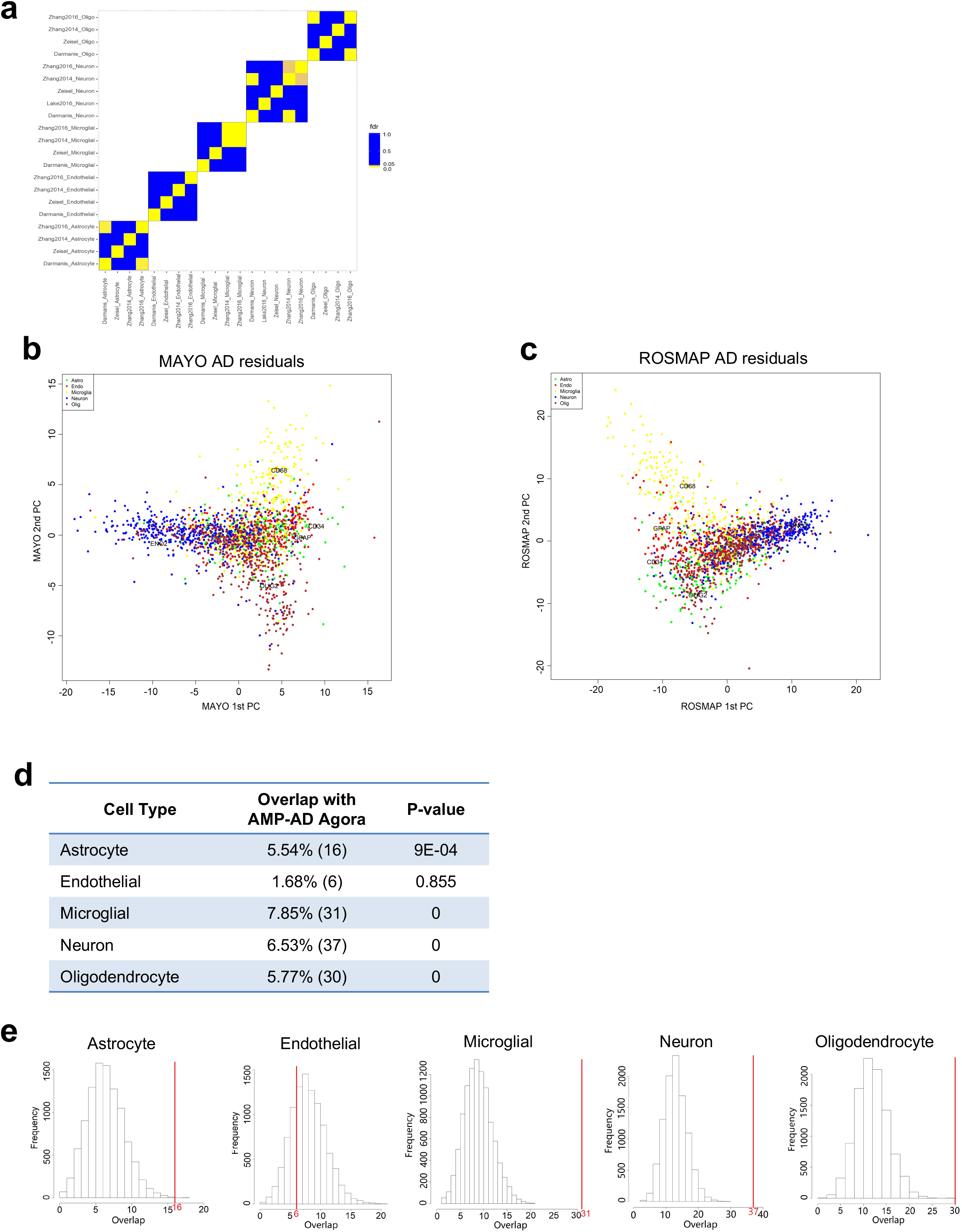
(**a**) Enrichment analysis of multiple multi-gene biomarker lists for the five main CNS cell types derived from various scRNAseq studies in control human brains shows no significant overlap among the studies. Significance was assessed by Fisher’s exact test with FDR cut-off of 0.05. (**b,c**) Principal component analysis (PCA) shows a prominent overlap of scRNAseq biomarker expression across the five main CNS cell types in AD residuals from the MAYO (**b**) and ROSMAP (**c**) datasets after covariate adjustment. (**d**) Enrichment analysis reveals significant overlap between scRNAseq biomarkers and AMP-AD Agora targets for neurons, microglia, astrocytes, and oligodendrocytes. The value in parentheses represents the number of genes overlapping between each biomarker list and the AMP-AD Agora targets. (**e**) Compared to randomly selected genes from the background overlapping with the AMP-AD Agora list, the number of overlapping genes between scRNAseq biomarkers and AMP-AD Agora targets (red vertical line along the distribution) is significantly higher for four cell types, including neurons. Significance was assessed by Fisher’s exact test with FDR cut-off of 0.05.

**Figure S4.**
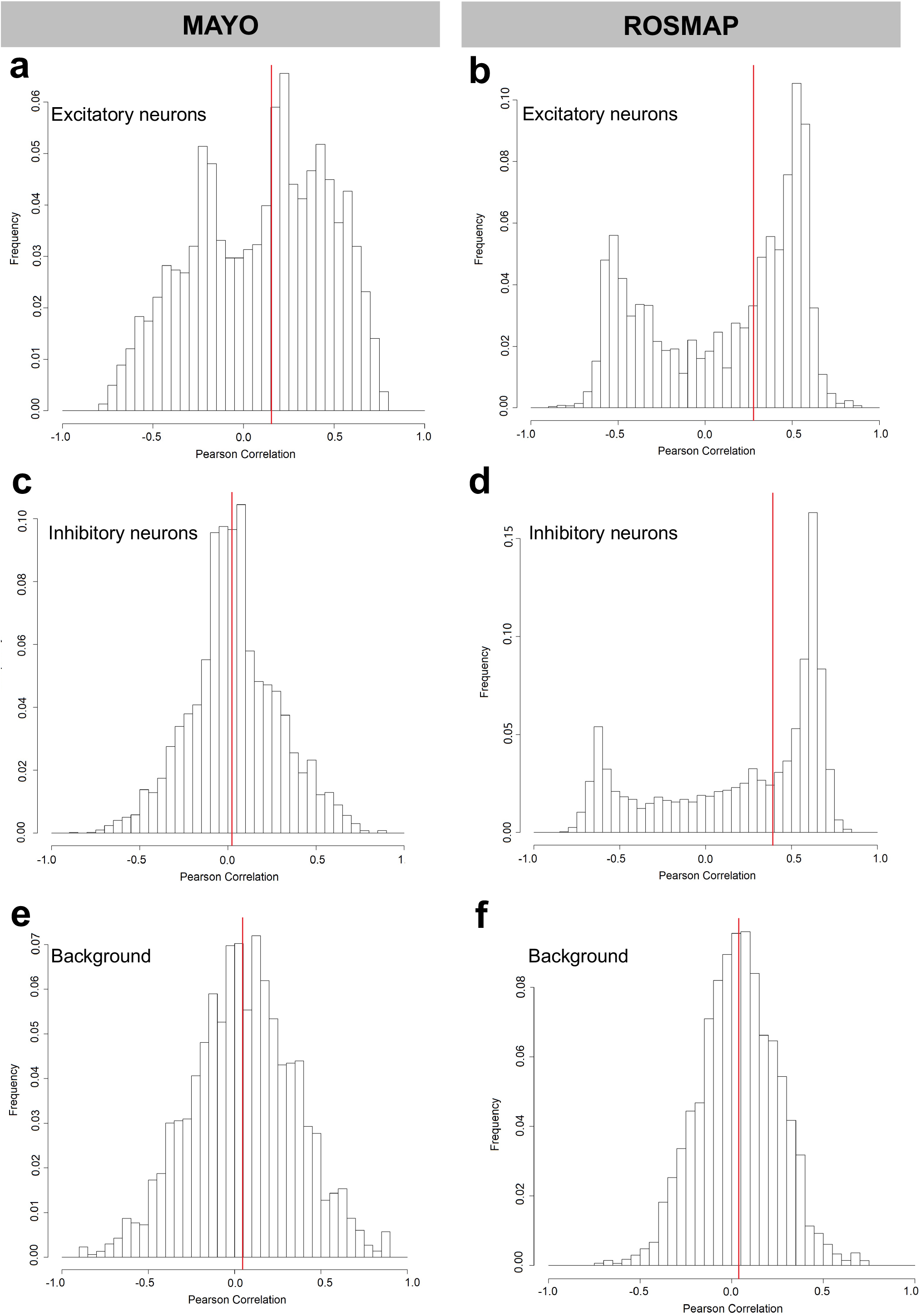
Distributions of Pearson correlation coefficients between our neuron-specific residual derived from *ENO2* and the “pseudo” neuron-specific residuals derived from a randomly selected subset of excitatory (**a,b**) and inhibitory (**c,d**) neuronal scRNAseq biomarkers by population-specific expression analysis (PSEA), for both the MAYO and ROSMAP datasets. (**e,f**) Distributions of Pearson correlation coefficients for both datasets between “pseudo” residuals derived from a randomly selected subset of background genes and “pseudo” neuron-specific residuals derived from a randomly selected subset of neuronal scRNAseq biomarkers. Red vertical line indicates the median of each distribution.

**Figure S5.**
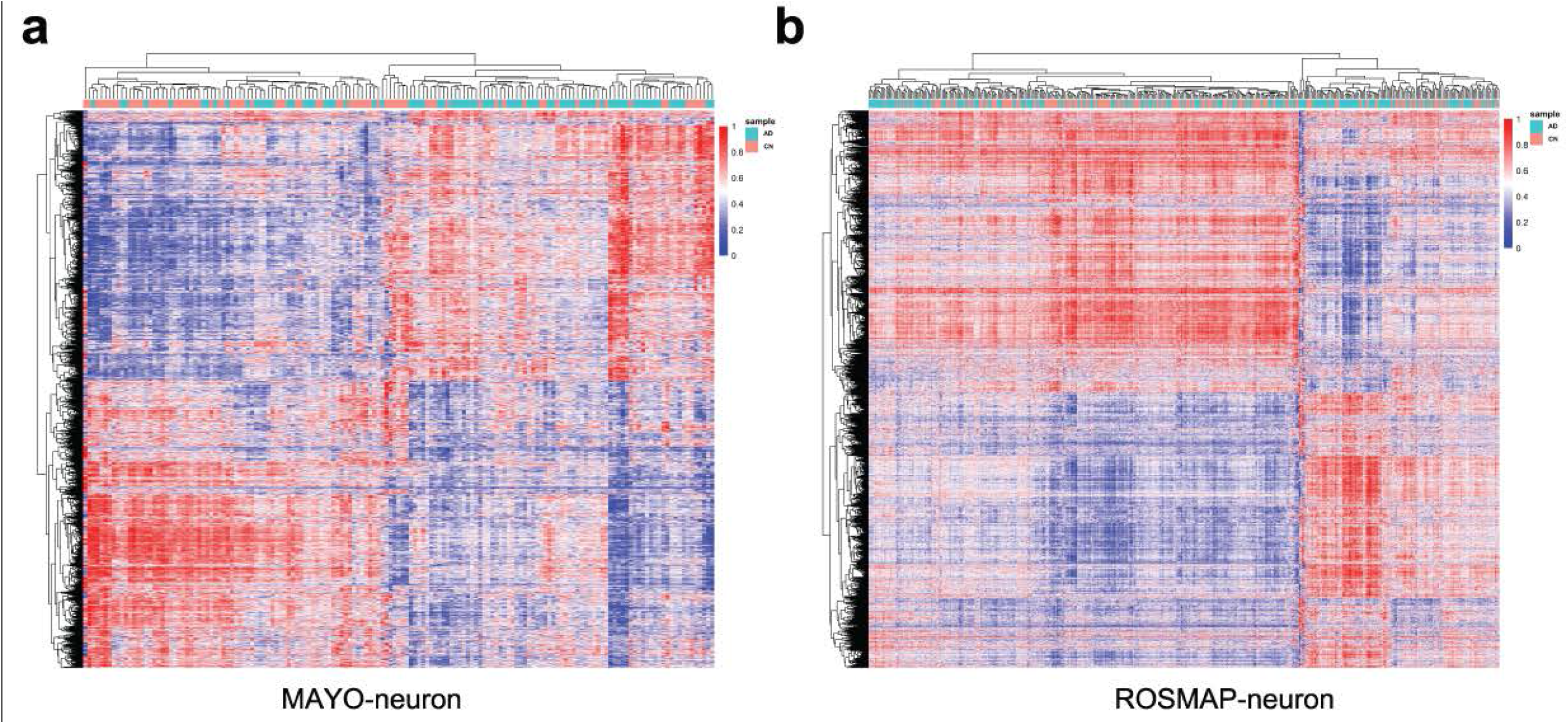
(**a,b**) Heatmaps showing clusters of upregulated (red) and downregulated (purple) genes in the neuron-specific residuals of AD patients compared to cognitively normal (CN) controls in MAYO (**a**) and ROSMAP (**b**).

**Figure S6.**
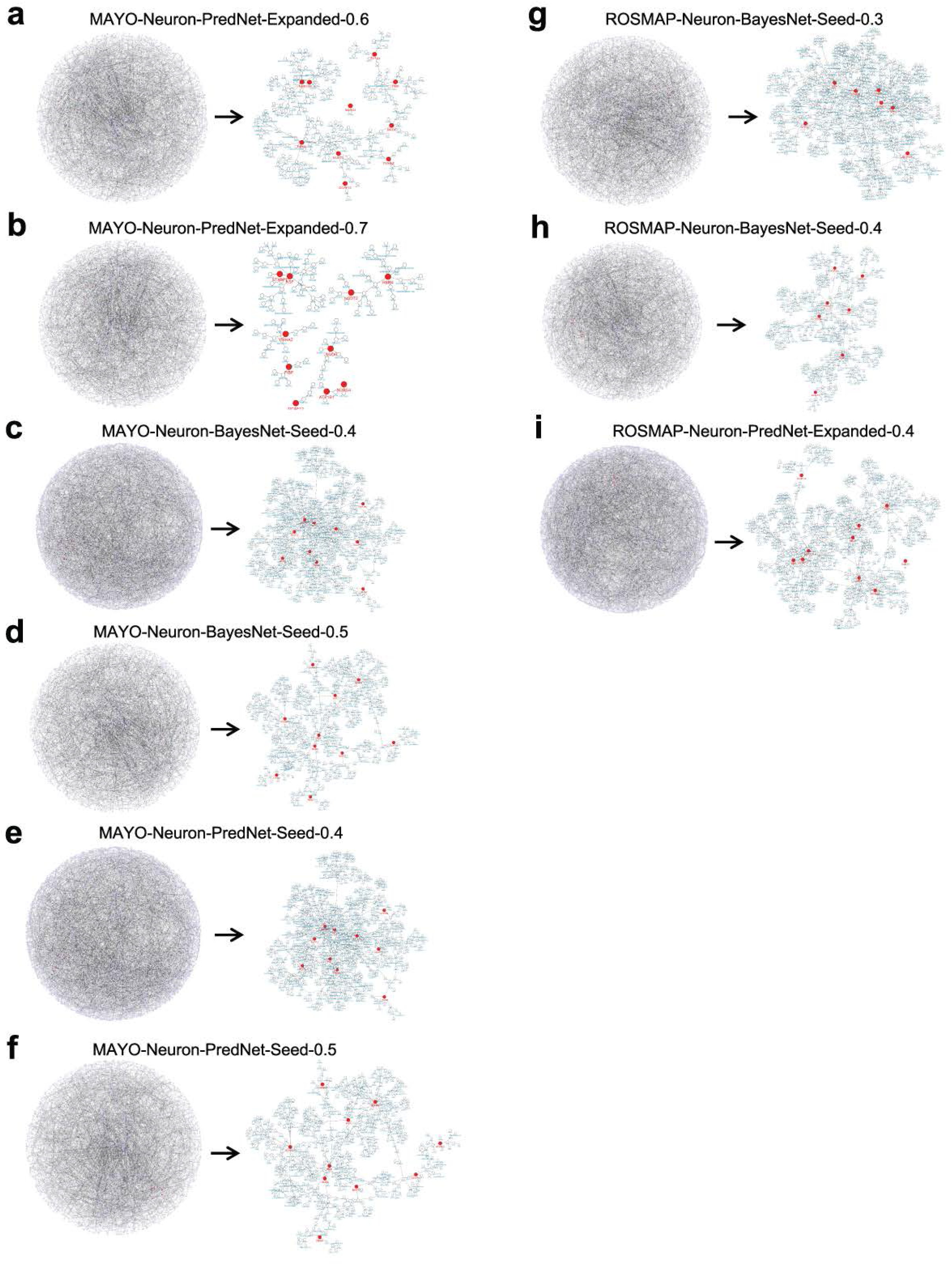
The remaining 9 of 11 neuron-specific Bayesian and predictive network models derived from the MAYO and ROSMAP seeding and expanded gene sets, with the downstream subnetworks of the 10 validated targets highlighted. Posterior probability cut-offs used to build each network model are indicated in each title.

**Figure S7.**
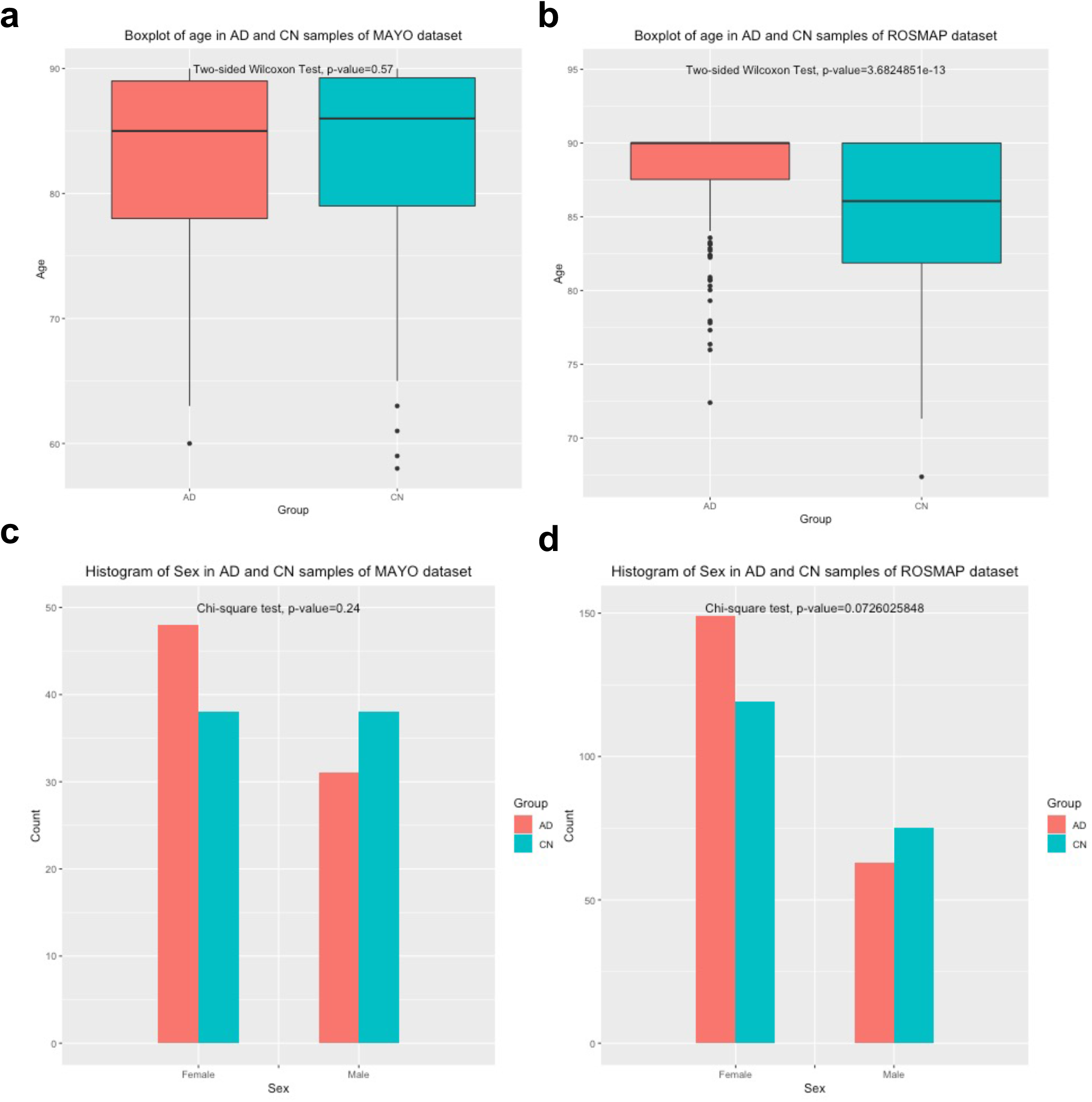
(**a,b**) The age distribution of AD and cognitively normal (CN) samples in the MAYO (**a**) and ROSMAP (**b**) datasets, each compared using an unpaired, two-sided Wilcoxon test. There was no significant age difference in MAYO (p-value=0.57) and a significant difference in ROSMAP (p-value=3.68E-13). To remove this effect in ROSMAP, age was adjusted along with other covariates in the ROSMAP residuals. (**c,d**) The sex distribution of AD and CN samples in the MAYO (**c**) and ROSMAP (**d**) datasets. A Chi-square test showed no significant difference in the sex breakdown in the MAYO (p-value=0.24) or ROSMAP (p-value=0.0726) datasets.

**Figure S8.**
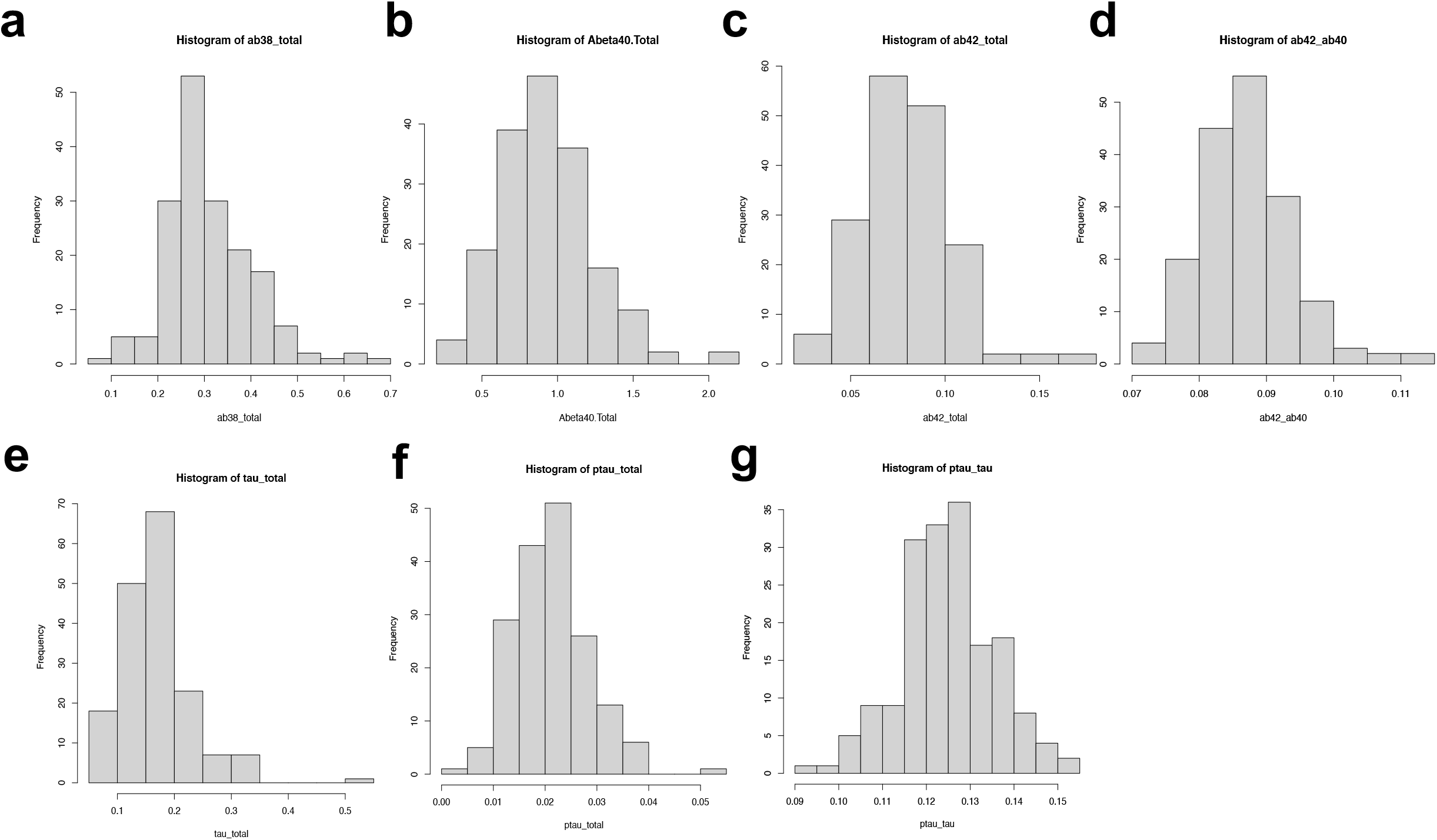
The frequency distributions of measurements for each AD endophenotype (Aβ38, Aβ40, Aβ42, Aβ42:Aβ40, p231-tau, tau, p231-tau:tau) found by pooling the values from all 19 shRNA targets plus controls for each parameter. These plots demonstrate normal and normal-like distributions for each measured AD endophenotype.

**Table S1.** Neuron-specific differentially expressed (DE) gene expression signatures associated with AD in the MAYO and ROSMAP RNAseq datasets. There are 2,097 significant DE genes overlapping between the two datasets.

**Table S2.** Significantly enriched pathways associated with neuron-specific DE gene signatures in the MAYO and ROSMAP RNAseq datasets, indicating dysregulated biological processes in AD. Pathway enrichment was assessed using Human ConsensusPathDB (CPDB).

**Table S3.** List of *cis*-eQTL genes in the MAYO and ROSMAP datasets.

**Table S4.** Significantly enriched biological pathways associated with gene modules in the MAYO and ROSMAP neuron-specific co-expression networks.

**Table S5.** Summary of statistical analyses of Aβ and tau data from human iNs following shRNA knockdown of each of the 19 prioritized key driver targets.

**Table S6.** Differential expression (DE) analysis of gene expression from RNAseq data from human iNs following shRNA knockdown of each of the 10 AD endophenotype modulating key driver targets.

**Table S7.** Significantly perturbed pathways associated with DE signatures of each of the 10 AD endophenotype modulating targets in human iNs.

**Table S8.** Cell-type specific biomarkers derived from existing scRNAseq studies for neurons, microglia, astrocytes, endothelial cells, and oligodendrocytes.

